# Rotational asymmetry is required to position centrioles at the base of the primary cilium

**DOI:** 10.64898/2026.07.08.732882

**Authors:** Meriem Boumendjel, Griselda Wentzinger, Mohamed Bahida, Tamara Advedissian, Tania Joanet, Futura Gattobigio, Farida Begum, Nicolas Moisan, Mark van Breugel, Takashi Ochi, Juliette Azimzadeh

## Abstract

The polarization of motile cilia requires that the centrioles, from which cilia are formed, display rotational asymmetry. This property is manifested in the presence of asymmetrically distributed appendages and relies on evolutionary conserved mechanisms. These mechanisms are also at play in cells that form primary cilia despite the lack of ciliary motility and asymmetric centriole appendages in this context. Here, we find that a complex consisting of CCDC61, KIAA1328 (K1328), and centlein (CNTLN) contributes to the establishment of centriole rotational asymmetry. In cells with a primary cilium, this complex is required for assembling a linker that repositions the daughter centriole close to and orthogonal to the proximal end of the mother centriole/basal body. The CCDC61/K1328/CNTLN complex also triggers the asymmetric recruitment of pericentriolar matrix components around newly assembled centrioles, which likely facilitates the later attachment of the basal body-daughter centriole linker. Overall, our results establish that rotational asymmetry relies on the coordinated recruitment of asymmetric landmarks along centrioles and is necessary for positioning the centrioles in a configuration that is widely conserved in ciliated cells.

## Introduction

Primary cilia are non-motile cilia that play essential roles in development and homeostasis by enabling the detection of biochemical and mechanical signals (Mill *et al*, 2023). Cilia are assembled from centrioles, which are composed of triplets of microtubules (MTs) in a 9-fold symmetric arrangement. Centrioles are found within the centrosome in most animal cells. The centrosome contains two centrioles, called mother (MC) and daughter (DC) centrioles, which are surrounded by a pericentriolar matrix (PCM) and connected by a filamentous linker. During the induction of ciliogenesis, the MC, which in this context is referred to as the basal body (BB), docks to the plasma membrane through its distal region and nucleates the formation of the cilium.

In ciliated protists and animal multiciliated cells (MCCs), centrioles carry appendages on specific MT triplets, rendering the centrioles rotationally asymmetric. These appendages are necessary to position the centrioles and control the direction of ciliary beating. Directional beating is essential for the locomotion or feeding of many protists, and for moving fluids such as respiratory mucus or cerebrospinal fluid at the surface of MCCs (Zhu, 2025). VFL1 and VFL3 are evolutionarily conserved proteins that are required for specifying the MT triplets on which asymmetric appendages are assembled. Inactivation of VFL1 and VFL3 leads to defects in asymmetric appendage assembly, which in turn cause defects in centriole position and ciliary beat orientation (Silflow *et al*, 2001; Bengueddach *et al*, 2017; Basquin *et al*, 2019; Nommick *et al*, 2022; Ochi *et al*, 2020; Wright *et al*, 1983; Adams *et al*, 1985). In contrast, the centrioles of the centrosome, from which primary cilia are assembled, do not carry asymmetric appendages. However, the molecular mechanisms governing centriole rotational asymmetry in animal MCCs and in protists appear to have been conserved at the centrosome. The human VFL1 ortholog, named LRRCC1, is enriched near two consecutive MT triplets in the distal region of centrioles in human cells, which is reminiscent of Vfl1p localization in the unicellular green alga *Chlamydomonas reinhardtii* (Gaudin *et al*, 2022; Silflow *et al*, 2001). LRRCC1 is involved in the organization of the distal end of the centriole and the assembly of the primary cilium, but why its asymmetric localization is conserved in the centrosome is unclear (Gaudin *et al*, 2022). In addition to the centrioles themselves, the distribution of the inner PCM layer, or torus, around the centrioles was recently found to be asymmetric in RPE1 cells, a non-transformed human cell line that forms primary cilia upon serum starvation. In these cells, the distribution of the torus component CEP152 and its partners CEP63 and CEP57 is most often interrupted by a gap spanning two or three consecutive triplets. In contrast, transformed cell lines such as HeLa and U2-OS, which can no longer form primary cilia, do not exhibit PCM asymmetry (Sullenberger *et al*, 2023).

The human VFL3 ortholog, called CCDC61, is conserved at the centrosome. CCDC61 interacts with CEP170 and ninein (NIN), two proteins that help to anchor MTs at the centrosome (Pizon *et al*, 2020; Barenz *et al*, 2018; Ochi *et al*, 2020; Carden *et al*, 2023). Like CEP170 and NIN, CCDC61 localizes to subdistal appendages (SDAs), which decorate the MC in a symmetric fashion and are involved in MT anchorage. In addition, all three proteins localize to the proximal end of centrioles, but their role there is unclear (Guarguaglini *et al*, 2005; Mogensen *et al*, 2000; Pizon *et al*, 2020). We have previously shown that CCDC61 contributes to the cohesion between centrioles (Pizon *et al*, 2020). Centriole cohesion depends on a linker formed by striated filaments containing the protein rootletin and several additional proteins. The rootletin linker is established at the beginning of the G1 phase of the cell cycle and degraded in G2 to allow the separation of duplicated centrosomes and mitotic spindle formation (Dang & Schiebel, 2022). In addition to rootletin filaments, MTs anchored to SDAs and NIN at the proximal ends of centrioles also contribute to centriole cohesion (Panic *et al*, 2015; Mazo *et al*, 2016; Theile *et al*, 2023). In RPE1, CCDC61 depletion enhances the splitting phenotype caused by inactivation of C-Nap1, a protein that anchors both rootletin and NIN to centriole proximal ends (Pizon *et al*, 2020). This suggests that CCDC61 contributes to centriole cohesion via its localization to SDAs. However, like NIN, CCDC61 could also participate in centriole cohesion through its proximal localization.

Here we show using expansion microscopy that CCDC61 localizes in a rotationally asymmetric manner along the proximal half of centrioles. We identify two novel CCDC61 interactors, KIAA1328 (K1328), also known as hinderin, and centlein (CNTLN), which both colocalize with CCDC61 in the proximal part of centrioles. The CCDC61/K1328/CNTLN complex is located on the same triplets as LRRCC1, but at opposite ends of the centriole. CCDC61 interacts and transiently co-localizes with LRRCC1 during centriole assembly to allow its recruitment to the same triplets. CCDC61 then recruits K1328 and CNTLN to centriole proximal end. The overall pattern of centriole rotational asymmetry defined by the four proteins is conserved in MCCs. We further demonstrate that centriole rotational asymmetry triggers PCM asymmetry in RPE1 cells. Our data support that the CCDC61/K1328/CNTLN complex is required for anchoring a linker, distinct from the rootletin linker, that connects the proximal end of the BB to the lateral side of the DC in cells with a primary cilium. This BB-DC linker contains CEP170 and NIN, but no rootletin, confirming that it is distinct from the rootletin linker.

## Results

### CCDC61 localizes asymmetrically at the proximal end of centrioles

We have previously determined the localization of CCDC61 in RPE1 cells by immuno-EM. These analyses revealed the presence of CCDC61 at SDAs, as well as less intense labeling at the proximal end of centrioles (Pizon *et al*, 2020). Because our anti-CCDC61 antibody did not work in immuno-EM, we used a cell line overexpressing a Myc-tagged version of CCDC61 approximately ten times the endogenous level. Immunofluorescence analysis of the endogenous protein also supported dual localization to SDAs and to the proximal end of centrioles, but with limited resolution. We thus turned to ultrastructure expansion microscopy (U-ExM) (Gambarotto *et al*, 2019) to better characterize the localization of endogenous CCDC61. Using this approach, CCDC61 was detected at the proximal part of the centrioles in WT RPE1 cells, but not at SDAs (Figure 1A). Strikingly, CCDC61 labeled the proximal half of centrioles in a rotationally asymmetric manner. Transverse views showed that CCDC61 was most often associated with 2 (57 % of centrioles) or 3 (28 %) consecutive triplets (Figure 1A, B). Co-labeling with the SDA component centriolin to distinguish centrioles showed that these values were similar between MC and DC, indicating that the number of labelled triplets does not vary during centriole maturation (Figure 1B, Supplemental Figure S1). An average 3D view obtained from 43 individual centrioles confirmed that CCDC61 was mainly associated with 2 consecutive triplets, in the proximal part of the triplets and at their cytoplasmic surface (Figure 1C). A similar pattern was observed when DC and MC images were averaged separately, confirming that this pattern does not depend on centriole age. We compared CCDC61 distribution along the centrioles to that of previously characterized centriole markers (Laporte *et al*, 2024) to determine if CCDC61 is associated with a particular subdomain. CCDC61 extended beyond the region corresponding to the cartwheel marked by SAS-6 in the procentrioles (Figure 1D) and overlapped with the inner centriole scaffold stained with Centrin2 (Le Guennec *et al*, 2020). However, CCDC61 distribution along the triplets was most similar to that of the torus component CEP152.

**Figure 1:**
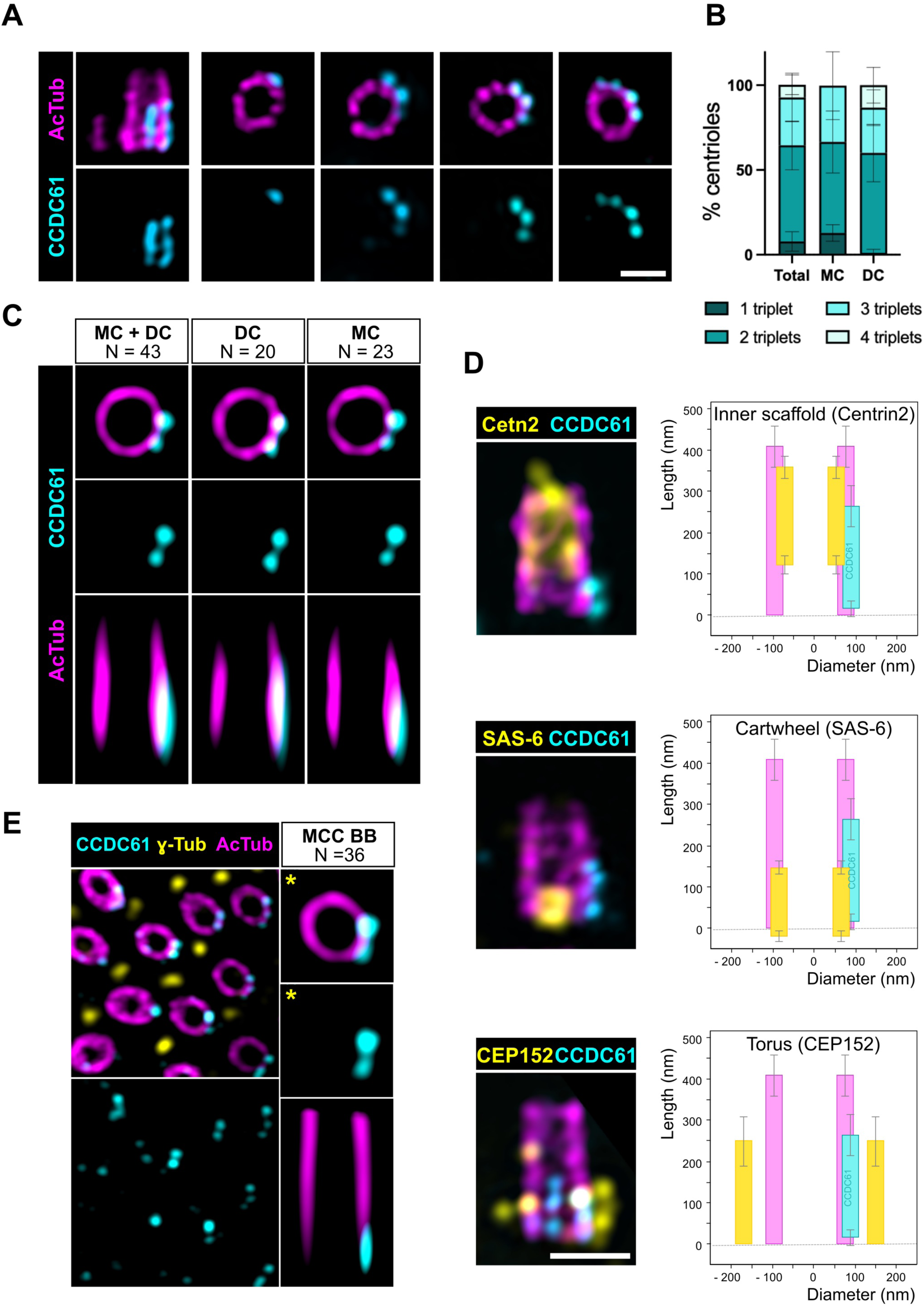
CCDC61 is located in a rotationally asymmetric manner in centrioles. **A)** U-ExM views of centrioles from WT RPE1 cells in longitudinal (left images) or transverse orientation. CCDC61 is shown in cyan and acetylated tubulin in magenta. Scale bar, 200 nm. **B)** Percentage of centrioles with CCDC61 labeling on 1, 2, 3 or 4 consecutive triplets in mother centrioles (MC), daughter centrioles (DC), or the combination of both (total). MC and DC were differentiated by staining the SDA component centriolin (Supplemental Figure S1). Four independent experiments, N = 150 centrioles. Bars represent the standard deviation. **C)** Average images obtained from 43 individual centrioles in transverse and longitudinal views (MC + DC). Average images obtained with only DCs or only MCs are also shown. CCDC61 is in cyan and acetylated tubulin is in magenta. **D)** Colocalization of CCDC61, in cyan, and markers of the inner scaffold (Centrin2), cartwheel (SAS-6) and inner PCM/torus (CEP152), in yellow, in RPE1 centrioles (acetylated tubulin, in magenta). The graphs show the average distribution of the different proteins relative to the proximal end (length) and the central axis (diameter) of the centriole. N = 42 (SAS-6), 46 (Centrin2, CEP152) and 213 (CCDC61) centrioles. Bars represent the mean and standard deviation. Note that Centrin2 is also located in the distal lumen of centrioles but this has not been included in the graph. **E)** Left, U-ExM view of a multiciliated cell (MCC) from the mouse trachea labeled with CCDC61 (cyan), γ-tubulin (yellow, basal foot marker), and acetylated tubulin (magenta). Right, average image obtained from 36 individual BBs, aligned with respect to the position of the basal foot (indicated by an asterisk).

The rotationally asymmetric localization of LRRCC1 in the centrioles of the centrosome is similar to that observed at the base of motile cilia in MCCs (Gaudin *et al*, 2022). To determine whether this is also true for CCDC61, we analyzed CCDC61 localization in mouse tracheal MCCs. The centrioles in MCCs show a conspicuous rotational asymmetry, manifested by the presence of the basal foot, an appendage that points in the direction of the ciliary beating (Lemullois *et al*, 1988). We thus determined CCDC61 localization with respect to the position of the basal foot labeled with anti-γ-tubulin (Figure 1E). Like in the centrosome, CCDC61 was most often localized on two consecutive triplets in the proximal part of the centrioles. These triplets were positioned opposite the basal foot, as also observed for LRRCC1 (Gaudin *et al*, 2022).

Taken together, these results show that CCDC61 is localized in a rotationally asymmetric manner, most often associating with two consecutive triplets in the proximal part of the centrioles, both at the centrosome and in MCCs.

### CCDC61 and LRRCC1 are located on the same MT triplets

Although CCDC61 and LRRCC1 are found at opposite ends of the centrioles, their localization with respect to the basal foot in MCCs suggested that they associate to the same triplets. Simultaneous labelling of CCDC61 and LRRCC1 in mouse tracheal MCCs confirmed that this is the case (Figure 2A). Similar analysis in RPE1 cells showed that CCDC61 and LRRCC1 also localize to the same triplets at the centrosome (Figure 2B). Thus, CCDC61 and LRRCC1 are localized in a coordinated manner, even though they are spatially segregated in full-length centrioles. To determine how their respective positions are coordinated, we analyzed CCDC61 and LRRCC1 recruitment during centriole assembly. We synchronized RPE1 cells using a double thymidine block and analyzed procentrioles by U-ExM at the end of the second block by labeling CCDC61, LRRCC1, and acetylated tubulin. A first category of procentrioles (7/176) had neither CCDC61 nor LRRCC1 and an average length of 80 nm based on acetylated tubulin labeling. This corresponds to an estimated total length of 123 nm, taking into account that the distal end of MT triplets is not acetylated at these stages (Figure 2C and Supplemental Figure S1 for the correlation between acetylated tubulin and total α/β-tubulin staining). This value is similar to that determined for the transition between the ‘bloom phase’, during which the triplets first assemble, and the elongation phase (Laporte *et al*, 2024). A second category (59/176) was positive for LRRCC1 only and corresponded to a later step of assembly (average length 146 nm). The last two categories contained both CCDC61 and LRRCC1, either colocalized at the distal end of the procentriole (57/176 procentrioles, average length 163 nm) or separated as in full length centrioles (53/176 procentrioles, average length 214 nm). These results indicate that LRRCC1 is recruited first at an early stage of procentriole assembly. CCDC61 is then recruited at the same location as LRRCC1, before the two proteins are separated by procentriole elongation.

**Figure 2:**
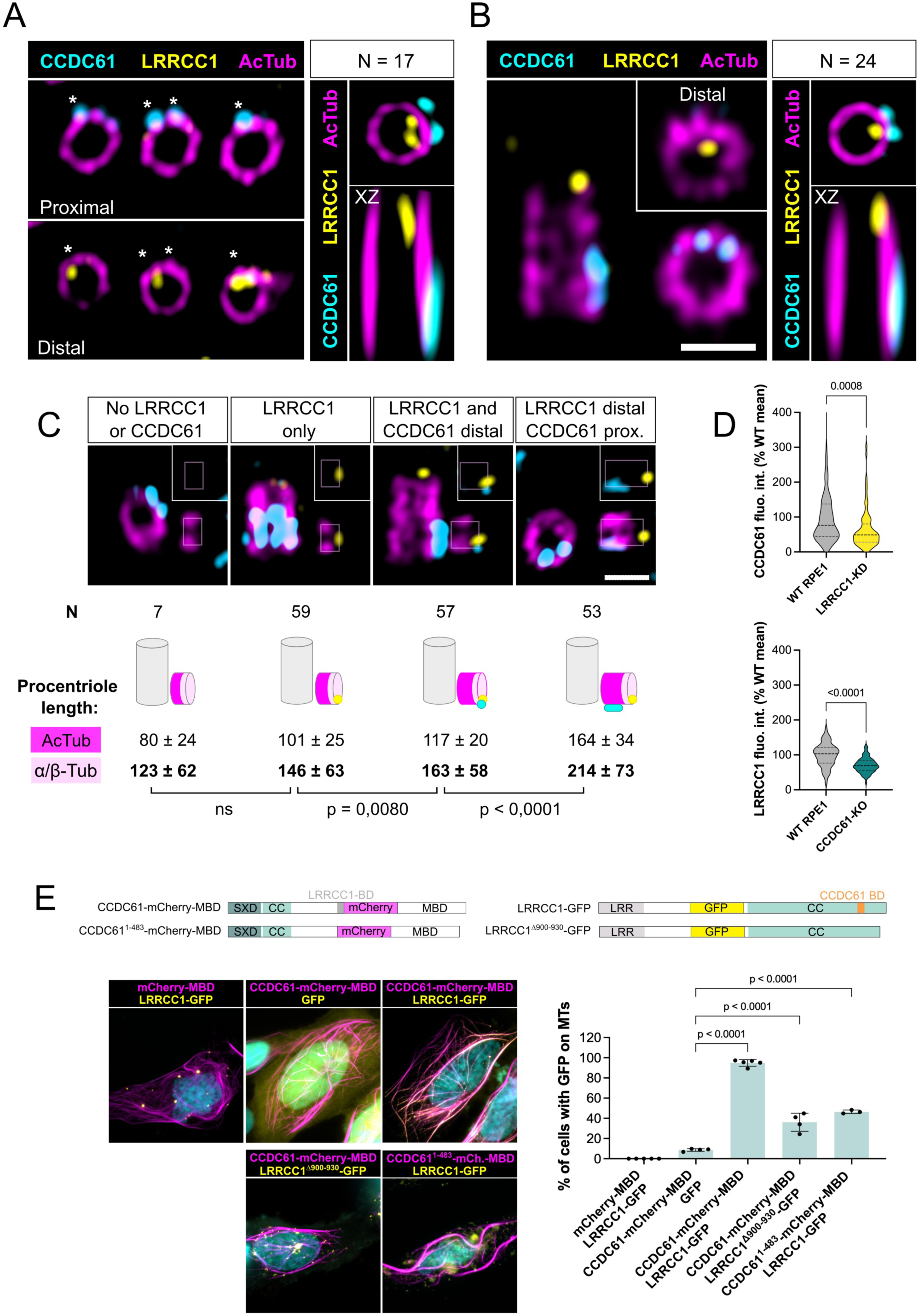
CCDC61 interacts with LRRCC1 and is located on the same triplets. **A)** Left panel: U-ExM view of individual centrioles in a mouse tracheal MCC co-stained for CCDC61 (cyan) and LRRCC1 (yellow), at the proximal or distal level. Acetylated tubulin in magenta. Right panel: average image obtained from 17 MCC centrioles co-stained for CCDC61 and LRRCC1, in transverse or longitudinal orientation. **B)** Left panel: U-ExM view of centrioles in a WT RPE1 cell co-stained for CCDC61 (cyan) and LRRCC1 (yellow). The centriole in transverse orientation is seen at the proximal or distal level (inset). Acetylated tubulin in magenta. Scale bar, 200 nm. Right panel: average image obtained from 24 RPE1 centrioles co-stained for CCDC61 and LRRCC1, in transverse or longitudinal orientation. **C)** U-ExM views of centrioles undergoing duplication (acetylated tubulin, magenta) showing the procentriole in longitudinal orientation. The procentrioles were systematically imaged and sorted according to the presence or absence of LRRCC1 (yellow) and CCDC61 (cyan). The number (N) and mean length ± standard deviation of procentrioles are indicated for each category. Procentriole length was determined using acetylated tubulin staining. Since the distal end of procentrioles is not acetylated, the estimated triplet length is also indicated (for the correspondence between acetylated tubulin and α/β-tubulin, see Supplemental Figure S1). p values are indicated when statistically significant (one-way ANOVA). Scale bar, 200 nm. **D)** Quantification of CCDC61 and LRRCC1 at centrioles in RPE1 WT and LRRCC1-KD or CCDC61-KO cells, respectively. Values are expressed as percentages of the corresponding mean control value. N > 85 centrioles for each condition. p values are indicated (Two-tailed T-test). **E)** Top: constructs used for the MT targeting assay. The interaction domains predicted by AlphaFold3 are indicated. MBD: MT-binding domain of FAM161A used to target the constructs to MTs. CC: coiled-coil; SXD: domain shared with SAS-6 and XRCC4 (Ochi *et al*, 2020). LRR: leucine-rich repeat. Bottom left: U2-OS cells expressing CCDC61, LRRCC1 or control constructs. Bottom right: percentage of cells with GFP constructs targeted to MTs. Three independent experiments, N > 80 for each condition. Bars indicate standard deviation. P values are indicated when relevant (one-way ANOVA).

The timing of recruitment and transient co-localization of the two proteins suggested that LRRCC1 could contribute to the recruitment of CCDC61. In agreement, CCDC61 centriolar levels were decreased in RPE1 cells partially depleted from LRRCC1 (65 % of control CCDC61 levels in a hypomorph LRRCC1 CRISPR mutant clone treated with RNAi to further decrease LRRCC1 levels, expressing about 25 % of control LRRCC1 levels) (Gaudin *et al*, 2022)(Figure 2D). The localization pattern of CCDC61 was not affected in LRRCC1-depleted cells, possibly due to the remaining LRRCC1 levels. LRRCC1 was also decreased in CCDC61-KO cells (70 % of control LRRCC1 levels), suggesting that CCDC61 might in turn stabilize LRRCC1 association to centrioles.

The co-localization and mutual dependence for their recruitment suggested that CCDC61 and LRRCC1 could interact directly. To test this, we used a previously described MT-targeting assay (Le Guennec *et al*, 2020). As overexpressed CCDC61 is only partially targeted to cytoplasmic MTs, we fused it to the MT binding domain (MBD) of the centriole component FAM161A (Figure 2E). A LRRCC1-GFP fusion was strongly recruited to MTs when co-overexpressed with MBD-CCDC61-mCherry, but not with the MBD-mCherry control, confirming that LRRCC1 can interact with CCDC61. AlphaFold3 predicted an interaction between a shorth segment (aa 900-930) of LRRCC1 coiled-coil domain and the C-terminal helix of CCDC61 (aa 484-512). These domains are conserved at the amino acid level, and interactions via the same domains were predicted in distantly related species (Supplemental Figure S2). Deleting either of these domains in the human proteins strongly reduced the recruitment of LRRCC1 to MTs, confirming that they are involved in the interaction between CCDC61 and LRRCC1 (Figure 2E). However, the interaction was not completely abolished, suggesting that other regions of the two proteins are involved. Comparisons between different fragments of the two proteins indicated that the central regions of CCDC61 and LRRCC1 also participate in the interaction (Supplemental Figure S2).

Together, these data show that CCDC61 and LRRCC1 define a consistent pattern of rotational asymmetry along the centrioles both in the centrosome and in MCCs. LRRCC1 is recruited first and contributes to recruiting CCDC61 to the same triplets, likely via a direct interaction.

### CCDC61 forms a complex with K1328 and CNTLN at the proximal end of centrioles

To identify additional putative CCDC61 interactors, we compared the proteome of centrosomes purified from wild type RPE1 cells with that of CCDC61-KO cells. Centrosomes were purified using the CAPture protocol and their composition was determined by mass spectrometry (Carden *et al*, 2023). Two proteins were fully missing in centrosomes purified from CCDC61-KO cells (Figure 3A): KIAA1328 (K1328)/hinderin, a predicted conserved component of centrioles (Drew *et al*, 2017; Carden *et al*, 2023), and centlein (CNTLN), an interactor of centrosome linker components C-Nap1 and CEP68 (Fang *et al*, 2014). To determine whether K1328 and CNTLN are potential CCDC61 interactors, we used the same MT targeting assay as previously. GFP-K1328 was found in cytoplasmic aggregates when co-expressed with the MBD-mCherry control, but was fully recruited to MTs in the presence of MBD-CCDC61-mCherry, supporting that K1328 interacts with CCDC61 (Figure 3B). GFP-CNTLN was in part targeted to MTs in approximately 40% of cells co-transfected with MBD-mCherry (not shown), supporting that CNTLN can bind MTs as described in a previous report (Jing *et al*, 2016). We thus fused CNTLN to the MBD and tested its interaction with the other two proteins. GFP-CCDC61 and GFP-K1328 were recruited to MTs in 90 % or 100 % of cells co-transfected with MBD-mCherry-CNTLN, compared to 25% or 0% of cells co-transfected with the MBD-mCherry control, respectively. These results support that each component of the complex is capable of interacting with the two others. This finding was confirmed by Western blot (Figure 3C).

**Figure 3:**
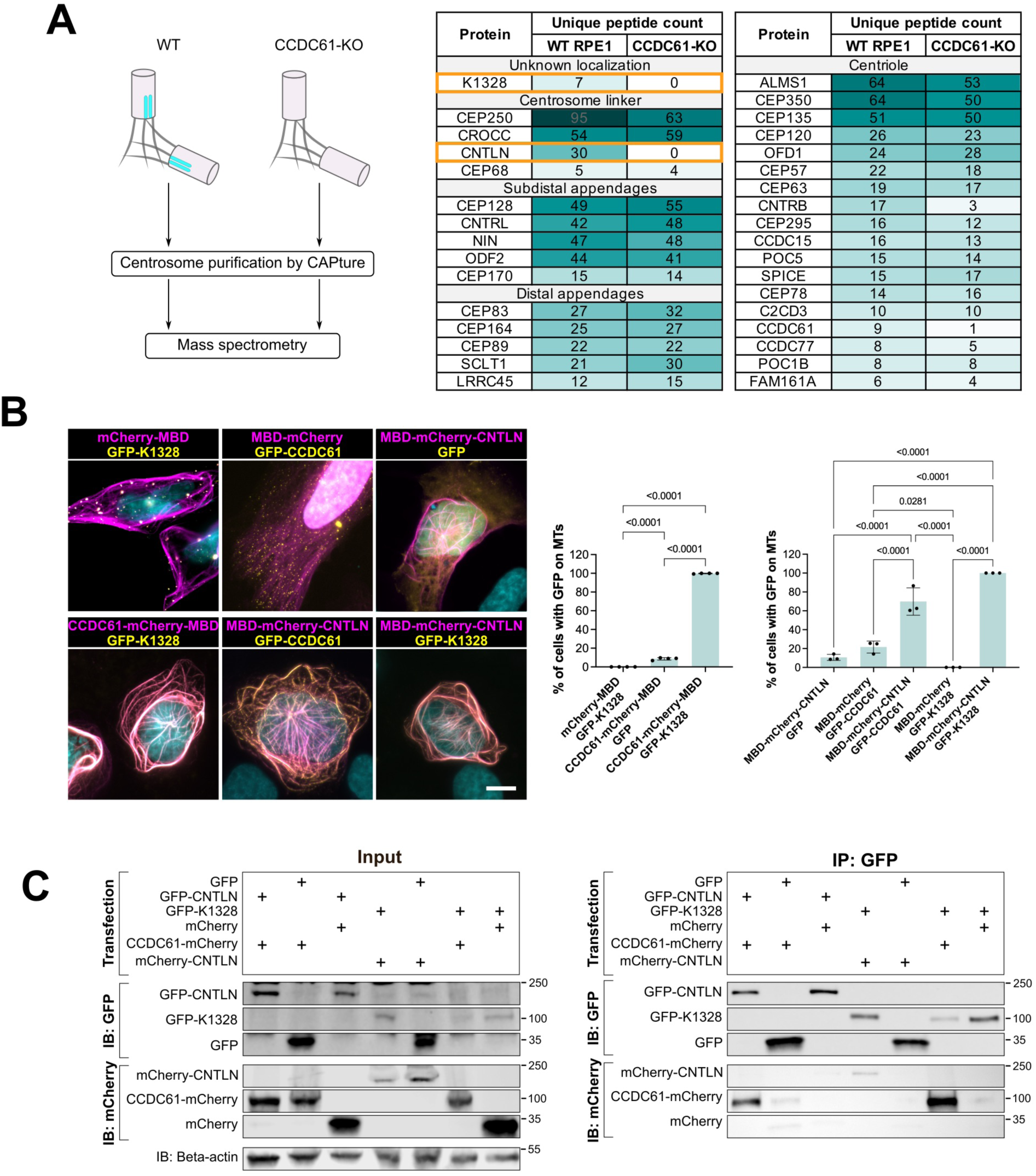
CCDC61 interacts with K1328 and CNTLN. **A)** Mass spectrometry identification of potential interactors of CCDC61. Centrosomes were purified by CAPture from RPE1 WT or CCDC61-KO cells and analyzed by mass spectrometry. The number of unique peptides identified is indicated for each protein. **B)** Left: U2-OS cells expressing GFP-K1328, GFP-CCDC61 or GFP with either the mCherry-MBD control construct, or MBD-mCherry-CNTLN. MBD: MT-binding domain of FAM161A. Right: percentage of cells with GFP constructs targeted to MTs. Three independent experiments, bars indicate standard deviation. p values are indicated when relevant (one-way ANOVA). **C)** Co-immunoprecipitation analysis of interactions between CCDC61, K1328, and CNTLN. HEK293T cells were transfected with the indicated combinations of constructs encoding the proteins tagged either with GFP or mCherry. Complexes were immunoprecipitated with GFP-trap beads (IP), and co-precipitated proteins were detected with anti-GFP or anti-mCherry (IB). β-actin was used as a loading control.

To determine whether K1328 and CNTLN co-localize with CCDC61 at the proximal end of centrioles, we examined their localization by U-ExM. In WT RPE1 cells, both proteins were mostly localized along two consecutive triplets in the proximal half of the centriole, a pattern highly similar to that of CCDC61 (Figure 4A, B). However, upon observing ciliated cells—in which the older centriole is easily identified by the presence of an associated cilium—we noticed that the staining was generally more intense on the DC. We measured the BB/DC ratio of the three proteins within individual centrosomes and found average values of 0.63 for CCDC61, 0.53 for K1328, and 0.48 for CNTLN (Figure 4C). Thus, even though the number of labeled triplets does not vary, the complex is approximately half as abundant on the BB as on the DC. Next, we examined whether the three proteins are colocalized. Simultaneous labeling confirmed that K1328 co-localize with CCDC61 in the proximal part of centrioles (Figure 4D). The CNTLN localization pattern also consistently mirrored that of CCDC61, but usually with a shift, suggesting that the epitopes recognized on the two proteins are distant from each other. Interestingly, the shift was always in the same direction, i.e. clockwise when the centrioles are viewed from the distal end (64/65 centrioles), suggesting these epitopes might be located near neighboring triplets. CNTLN was not detected at the proximal tip of centrioles or between MC and DC, as would be expected for a component of the rootletin and CEP68-containing linker. The amounts of both K1328 and CNTLN were strongly decreased in CCDC61-KO centrioles (16 % and 2 % of control levels, respectively), confirming that these proteins are recruited to centrioles in a CCDC61-dependent manner (Figure 4E). In a K1328-KO cell line obtained by CRISPR/Cas9 editing (Supplemental Figure S3), centriolar CNTLN was reduced to 12 % of control levels (Figure 4E). Conversely, CCDC61 was decreased to 40 % of control levels in K1328-KO centrioles, indicating that K1328 in turn contributes to recruiting or stabilizing CCDC61 at centrioles. We obtained two CRISPR-knockdown (KD) CNTLN cell lines that express mutant versions of the protein, with a residual staining at the centrosome corresponding to approximately 10% of the wild type levels (Supplemental Figure S3). In both cell lines, the levels of CCDC61 and K1328 were decreased compared to controls (approximately 70 % and 30-50 % of control levels, respectively), suggesting that CNTLN also helps recruiting or stabilizing these proteins. Overall, these data show that CCDC61 is required for the recruitment of K1328 to the centrioles, and that K1328 is in turn required for the recruitment of CNTLN. Conversely, K1328 and CNTLN contribute to the recruitment or stabilization of CCDC61 and K1328, respectively.

**Figure 4:**
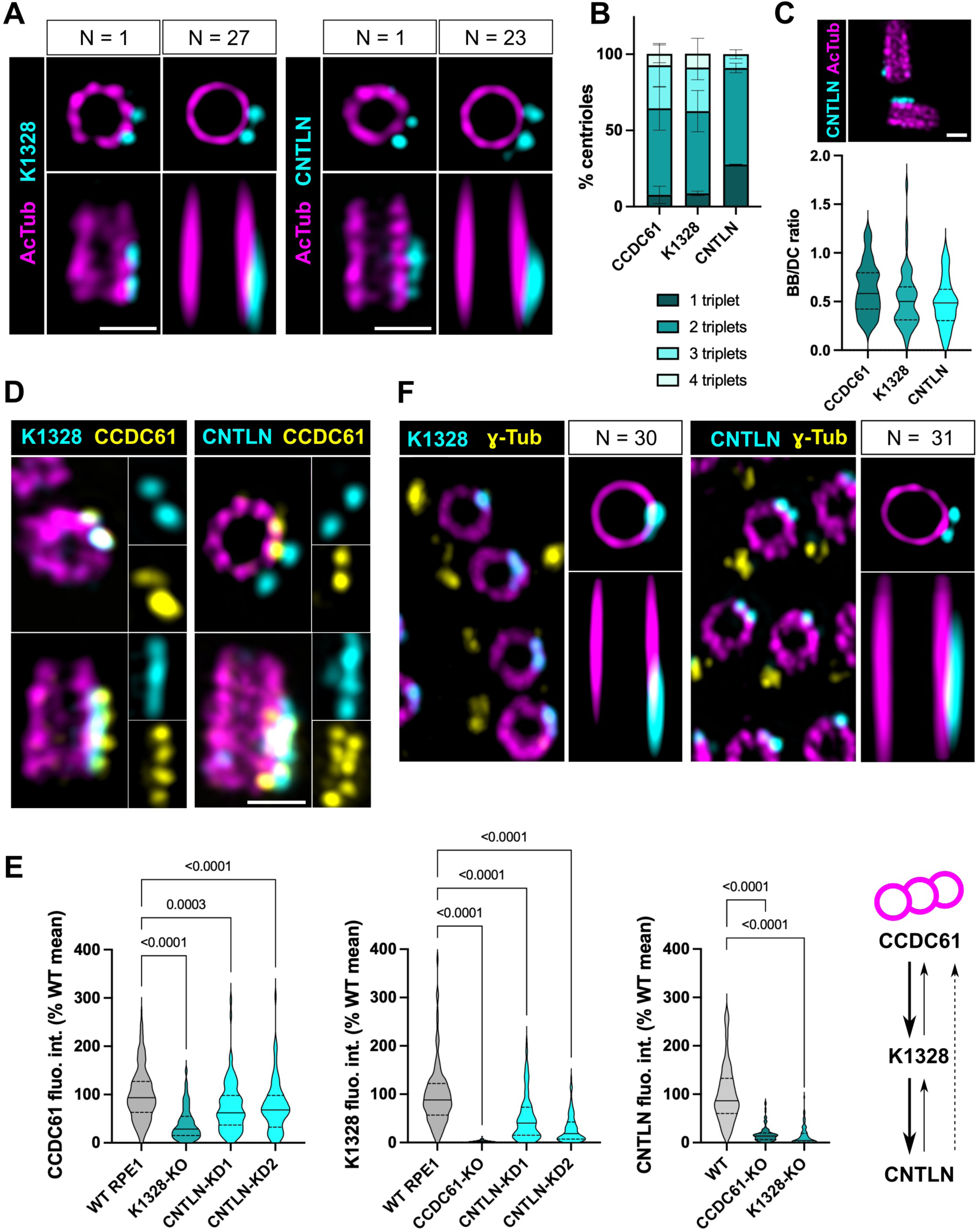
K1328 and CNTLN colocalize with CCDC61 at the proximal end of centrioles. **A)** U-ExM views of individual centrioles or average images in RPE1 cells stained with either K1328 or CNTLN (cyan) and acetylated tubulin (magenta). **B)** Percentage of centrioles with CCDC61 labeling on 1, 2, 3 or 4 consecutive triplets. Three independent experiments, N = 150 (CCDC61), 36 (K1328) and 53 (CNTLN) centrioles. Bars represent the standard deviation. **C)** Top: U-ExM view of a WT ciliated RPE1 cell stained for CNTLN (cyan) and acetylated tubulin (magenta). Bar, 200 nm. Bottom: BB/DC ratio of fluorescence intensity for CCDC61, K1328, and CNTLN in WT ciliated RPE1 cells. N = 45 (CCDC61), 51 (K1328) and 52 (CNTLN) centriole pairs. **D)** U-ExM images of RPE1 centrioles co-stained for CCDC61 (yellow) and either K1328 or CNTLN (cyan). Acetylated tubulin is in magenta. Scale bar, 200 nm. **E)** Quantification of CCDC61, K1328, and CNTLN in RPE1 WT, CCDC61-KO, and K1328-KO cells in U-ExM, as a percentage of the corresponding control mean. Three independent experiments, N> 70 centrioles for each condition. p values are indicated when the difference is significant (one-way ANOVA). Right: Hierarchical recruitment of CCDC61, K1328, and CNTLN based on these analyses. **F)** U-ExM views of mouse tracheal MCCs labeled with K1328 or CNTLN (cyan), γ-tubulin (yellow, basal foot marker), and acetylated tubulin (magenta). Right panels show average images in transverse or longitudinal orientation. Individual BBs were aligned with respect to the position of the basal foot.

We next examined the localization of K1328 and CNTLN in mouse tracheal MCCs. Both proteins had a localization pattern similar to that of CCDC61, i.e. mainly associated with two triplets opposite the basal foot (Figure 4F). This indicates that CCDC61, K1328 and CNTLN also colocalize at the base of motile cilia.

Together, these results show that CCDC61, K1328 and CNTLN form a complex located asymmetrically in the proximal half of centrioles.

### The CCDC61/K1328/CNTLN complex drives the asymmetric recruitment of PCM components in RPE1 cells

In RPE1 cells, the CEP57/CEP63/CEP152 complex is recruited in an asymmetric fashion around centrioles, leaving a gap corresponding to 2 to 3 consecutive triplets (Sullenberger *et al*, 2023). As these proteins localize in the proximal half of the centriole as CCDC61 and its partners, we wondered whether the triplets lacking the CEP57/CEP63/CEP152 complex might be the ones decorated by CCDC61, K1328 and CNTLN. Simultaneous labeling of CCDC61 and CEP152 showed that CCDC61 was either entirely (20/28 centrioles) or at least partially (8/28 centrioles) located at CEP152-deficient triplets when CEP152 distribution was asymmetric (Figure 5A). Since CEP152 is necessary for procentriole assembly, this should result in fewer procentrioles forming opposite CCDC61-labeled triplets. In agreement, virtually no procentrioles were observed near CCDC61 labelled triplets (Figure 5B). We then sought to determine how the recruitment of the two complexes is rendered mutually exclusive. CCDC61 is recruited to procentrioles in S phase, while CEP57 and downstream components CEP63 and CEP152 are recruited at the onset of the following cell cycle (Watanabe *et al*, 2019; Wei *et al*, 2020; Zhao *et al*, 2020; Sir *et al*, 2011; Laporte *et al*, 2024). CCDC61 and its partners could thus trigger the asymmetry in CEP57/CEP63/CEP152 localization. In WT RPE1 cells, CEP152 was most often (75 % of centrioles) localized in an asymmetric manner (Figure 5C), as previously shown (Sullenberger *et al*, 2023). In contrast, in ∼ 90 % of CCDC61-KO and K1328-KO centrioles, and ∼ 70 % of CNTLN-KD centrioles, CEP152 distribution was symmetric, supporting that the presence of the CCDC61/K1328/CNTLN complex inhibits CEP152 recruitment to the same triplets. In addition to CEP63 and CEP152, CEP57 is also required for the recruitment of pericentrin (PCNT), another major component of the PCM (Ito *et al*, 2021; Zhao *et al*, 2020; Watanabe *et al*, 2019; Wei *et al*, 2020). It was therefore plausible that PCNT would also show an asymmetric distribution in RPE1 centrioles. In agreement, analysis of PCNT in wild type RPE1 cells revealed a predominantly asymmetric distribution (Figure 5D). When the two proteins were co-labeled, CCDC61 was localized on triplets that lacked PCNT in all centrioles displaying asymmetric PCNT distribution (15/15 centrioles) (Figure 5E). As previously, inactivation of CCDC61, K1328 or CNTLN led to a drastic change in PCNT distribution, from asymmetric to symmetric (Figure 5D).

**Figure 5:**
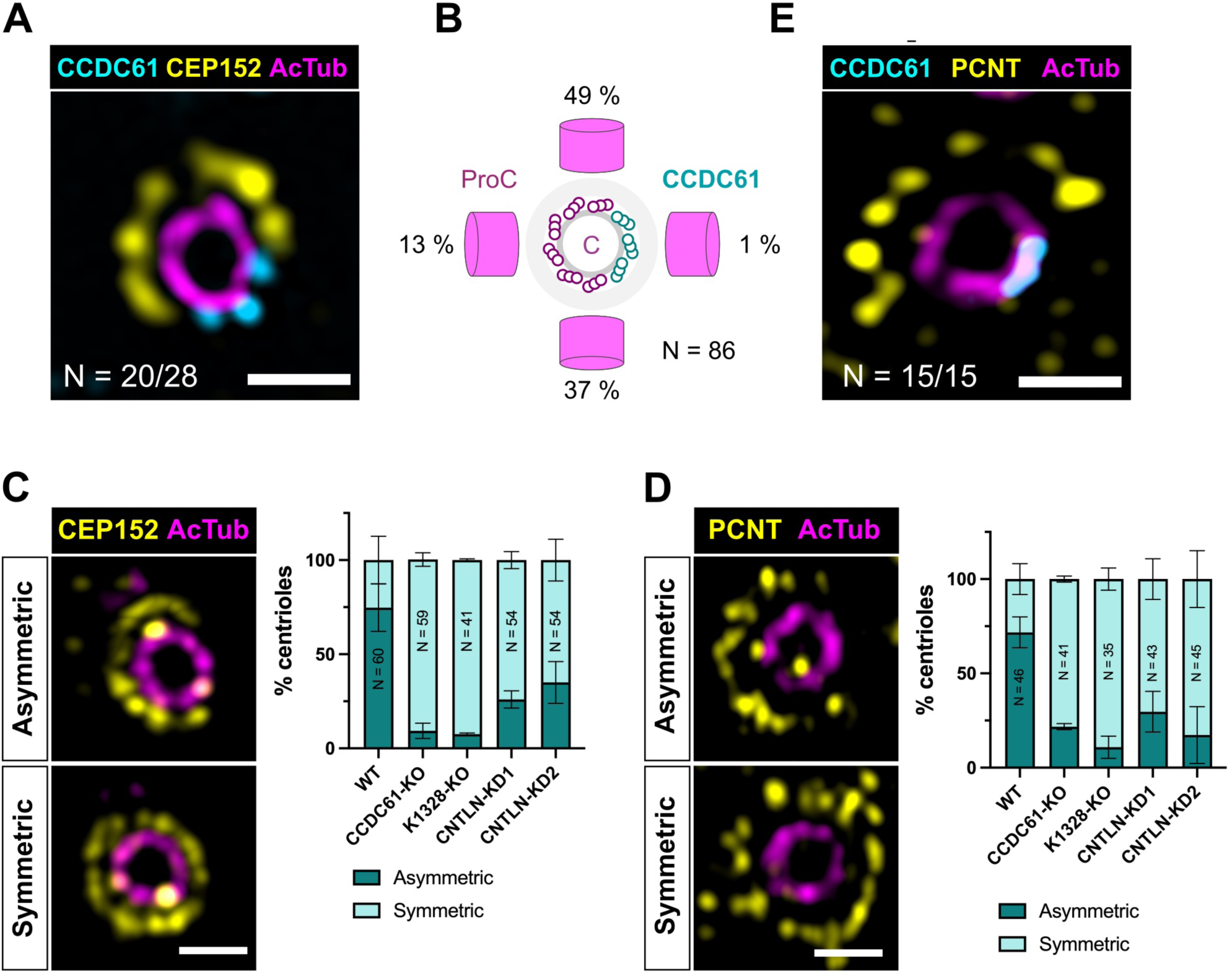
The CCDC61/K1328/CNTLN complex promotes asymmetric PCM recruitment in RPE1 cells. **A)** U-ExM view of an RPE1 centriole stained for CCDC61 (cyan), CEP152 (yellow), and acetylated tubulin (magenta). Bar, 200 nm. **B)** Schematic showing the frequency with which the procentriole (ProC) is formed relative to the position of CCDC61-labeled triplets on the parent centriole (C). The data were obtained from U-ExM images of WT RPE1 cells synchronized at the G1/S transition using thymidine. N = 86 procentrioles. **C)** Left: U-ExM images showing asymmetric or symmetric distribution of CEP152 (yellow) around WT RPE1 centrioles stained with acetylated tubulin (magenta). Bar, 200 nm. Right: frequency of asymmetric distribution of CEP152 in WT, CCDC61-KO, or K1328-KO RPE1 cells. 41 > N > 60 centrioles per condition, 2 to 4 independent experiments. Bars represent the mean and standard deviation. **D)** Left: U-ExM images showing asymmetric or symmetric distribution of PCNT (yellow) around WT RPE1 centrioles stained with acetylated tubulin (magenta). Bar, 200 nm. Right: frequency of asymmetric distribution of PCNT in WT, CCDC61-KO, or K1328-KO RPE1 cells. 35 > N > 46 centrioles per condition, 2 or 3 independent experiments. Bars represent the mean and standard deviation.

Collectively, these results demonstrate that asymmetric recruitment of the CCDC61/K1328/CNTLN complex is responsible for the asymmetric distribution of PCM components in RPE1 cells. The different landmarks of rotational asymmetry are thus interconnected, revealing a consistent pattern along the centrioles and in the PCM.

### The CCDC61/K1328/CNTLN complex is required for positioning the DC close to the BB in ciliated cells

We next investigated the functions associated with CCDC61/K1328/CNTLN asymmetric localization. We and others have shown that intercentriolar distance is decreased in ciliated cells, suggesting a remodeling of centriole linkage and a repositioning of the DC relative to the BB upon cilium formation (Pizon *et al*, 2020; Loukil *et al*, 2017). Our study further showed that inactivation of CCDC61 increased intercentriolar distance in ciliated RPE1 cells, indicating that DC repositioning is perturbed (Pizon *et al*, 2020). We observed a similar phenotype in ciliated K1328-KO and CNTLN-KD cells, supporting that K1328 and CNTLN contribute to DC repositioning together with CCDC61 (Figure 6A). We hypothesized that the CCDC61/K1328/CNTLN complex might enable a connection between centrioles specific to ciliated cells. We first conducted a detailed reassessment of the electron microscopy data available in the literature. We noticed that in ciliated cells from a variety of mammalian tissues, the linker between the BB and the DC, although at first glance resembling rootletin filaments, actually exhibit distinct characteristics. While the periodicity of the striations in rootletin filaments is approximately 75 nm, it is closer to 60 nm for the BB-DC linker. Furthermore, the BB-DC linker has a fixed pattern with three prominent transverse striations for a total length of approximately 250 nm, unlike rootletin filaments, which are longer and more variable in both length and diameter. Finally, the BB-DC linker connects the proximal end of the BB to the lateral side of the DC, to which it is attached by two or three consecutive triplets, while rootletin filaments are anchored to the proximal ends of both centrioles (Yamamoto & Kataoka, 1986; Hagiwara *et al*, 2002, 2008; Gallagher, 1980; Webber & Lee, 1975; Wheatley, 1967). Thus, the BB-DC linker has a distinct architecture from that of the rootletin linker, and the way it attaches to the side of the DC suggests that this may be mediated by the CCDC61/K1328/CNTLN complex. To determine whether this is the case, we first assessed the presence of the BB-DC linker in RPE1 cells. A previous observation by cryo-electron tomography of centrosomes purified from ciliated RPE1 cells suggested that this might be the case (Carden *et al*, 2023). Further analysis showed that when the connection between the DC and BB was maintained, the DC was positioned at the proximal end of the BB and perpendicular to it, consistent with the presence of the BB-DC linker. Moreover, the structure of the linker in RPE1 cells was similar to that observed by electron microscopy in earlier studies (Figure 6B). It consists of two layers of fine filaments approximately 60 nm long, oriented perpendicular to the DC. Electron-dense transverse striations delimit these layers (arrows in Figure 6B). The striation closest to the DC is approximately 40 nm away from the triplets. The striation closest to the BB is separated from it by a less well-defined region 100–150 nm wide.

**Figure 6:**
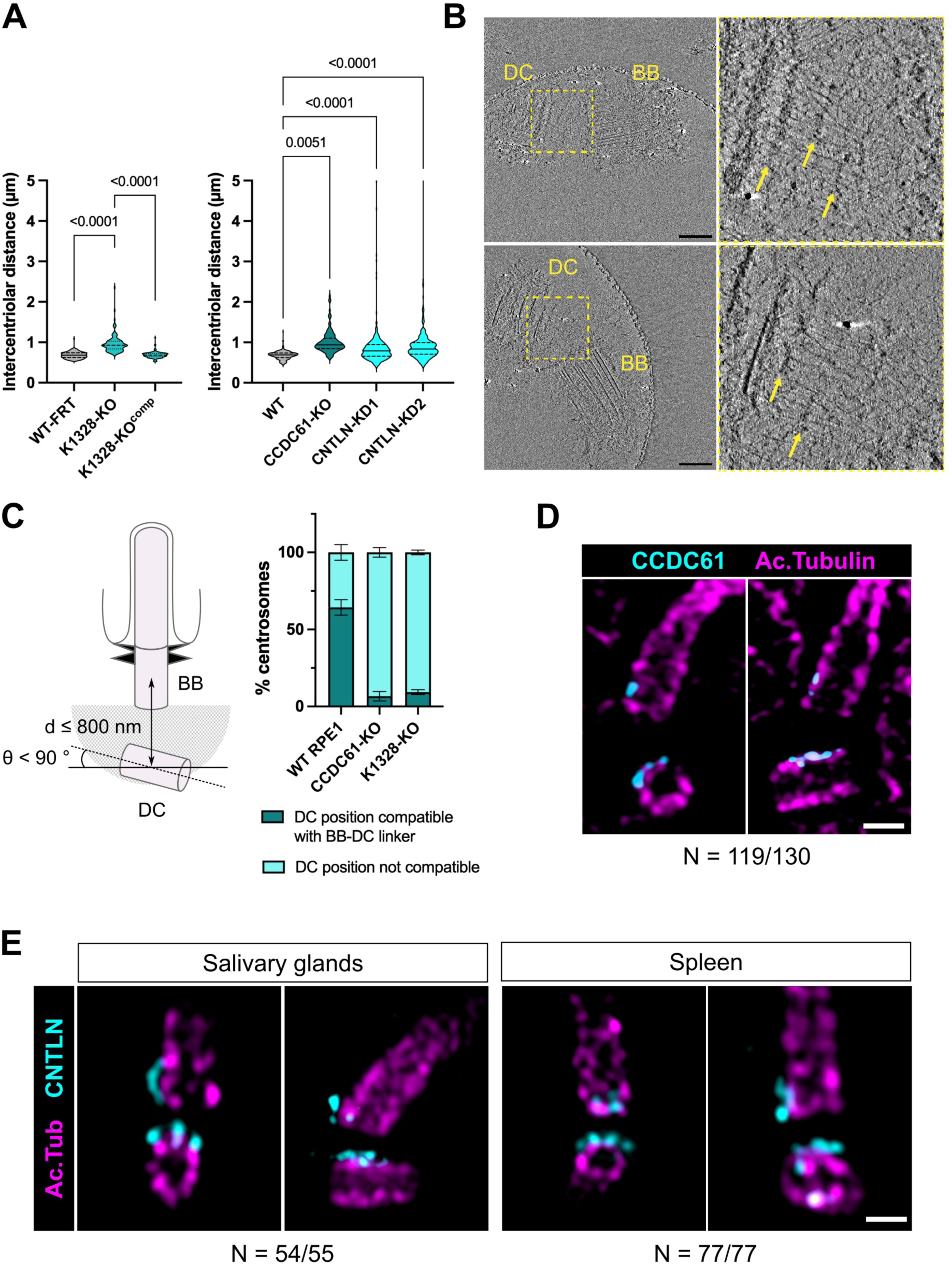
The CCDC61/K1328/CNTLN complex enables DC repositioning via the BB-DC linker in WT RPE1 cells. **A)** Intercentriolar distance 24 hours after induction of ciliogenesis by serum deprivation in K1328-KO, CCDC61-KO and CNTLN-KD cells. WT-FRT: RPE1 cells with a Flp-In recombination site, control for K1328-KO. K1328-KO^comp^: K1328-KO cells expressing K1328-GFP. Three independent experiments, 100 < N < 150 for each condition. p values are indicated when relevant (One-way ANOVA). **B)** Cryo-electron tomography of centriole pairs obtained by CAPture from serum-deprived RPE1 cells. The BB and DC are perpendicular to each other and connected by the BB-DC linker. The regions marked with a yellow rectangle in the left images are enlarged on the right. The arrows indicate the main striations of the BB-DC linker. Scale bar, 200 nm. **C)** Percentage of cells in which the position of the DC relative to the BB is compatible with the presence of a BB-DC linker. The criteria used are the distance between centrioles (d ≤ 800 nm between DC and BB centroids) and DC position (DC present in the proximal region of the BB and oriented sideways). Three independent experiments were performed, N > 300 cells for each condition. Bars represent the standard deviation. **D)** CCDC61-labeled triplets (cyan) on the DC are oriented toward the proximal end of the BB in the majority of WT ciliated RPE1 cells in which the position of the DC is compatible with the presence of the BB-DC linker. Acetylated tubulin is shown in magenta. Scale bar, 200 nm. **E)** The triplets marked with CNTLN (cyan) on the DC are oriented toward the proximal end of the BB in the majority of ciliated cells in mouse salivary glands and spleen. Acetylated tubulin is in magenta. Bar, 200 nm.

To estimate the frequency with which the BB-DC linker forms in RPE1 cells, we analyzed the respective positions of the DC and MC in ciliated RPE1 cells in U-ExM. In WT cells, the position of the DC was compatible with the presence of the BB-DC linker in terms of distance and orientation in around 65 % of cases (Figure 6C). This proportion fell to less than 10% for CCDC61-KO and K1328-KO cells. Thus, the BB-DC linker appears to be formed in a majority of RPE1 cells under our experimental conditions but is not present in CCDC61-KO and K1328-KO cells. We next determined whether the localization of the CCDC61/K1328/CNTLN complex on the DC in ciliated RPE1 cells was consistent with involvement in BB-DC linker attachment. In > 90 % of cases where the respective positions of the MC and DC indicated the presence of a BB-DC linker, CCDC61 was positioned on the side of the DC closest to the proximal end of the MC, where the BB-DC linker is anchored (Figure 6D). We then investigated whether this is also the case in mouse tissues. We focused on the spleen and salivary glands, for which we obtained the best image quality, and labeled CNTLN, which generates the strongest signal. In both tissues, the position of CNTLN on the DC was consistent with the presence of the BB-DC linker in virtually all cells analyzed (Figure 6E).

Together, these results support that repositioning of the DC close to the BB in ciliated cells involves a linker that specifically connects the proximal end of the BB to the lateral side of the DC via the CCDC61/K1328/CNTLN complex.

### CCDC61 interactors NIN and CEP170 are components of the BB-DC linker

Finally, we investigated the composition of the BB-DC linker. The different striation patterns suggested that the BB-DC linker has a distinct composition from that of rootletin filaments. Consistent with this, we most of the time (45/48 centrosomes) did not detect rootletin between centrioles in ciliated RPE1 cells in which DC position was indicative of the presence of the BB-DC linker (Figure 7A). CEP170 and NIN were good candidates to be components of the linker because they interact with CCDC61 and localize at the proximal end of the centrioles, where they are recruited by C-Nap1 (Mazo *et al*, 2016; Lamla, 2008; Barenz *et al*, 2018; Ochi *et al*, 2020; Carden *et al*, 2023). Furthermore, proximal NIN has been shown to contribute to centriole cohesion in RPE1 cells (Theile *et al*, 2023), and the increase in intercentriolar distance induced by NIN RNAi in ciliated RPE1 cells is comparable to what we observed in CCDC61-KO, K1328-KO and CNTLN-KD (Supplemental Figure S4). One possibility, therefore, was that the NIN and CEP170 fractions at the proximal end of the BB may contribute to the formation of the BB-DC linker by interacting with the CCDC61/K1328/CNTLN complex on the DC side. In the majority of ciliated RPE1 with properly positioned centrioles, NIN and CEP170 formed a linear pattern in the region between BB and DC that ran parallel to the CCDC61-labeled triplets on the DC side, as expected for putative components of the BB-DC linker (Figure 7B). In contrast, NIN and CEP170 were not recruited along the DC side in CCDC61-KO or K1328-KO ciliated cells, supporting that this pattern is dependent on the presence of the CCDC61/K1328/CNTLN complex on the DC. Interestingly, when the two centrioles were close together in CCDC61-KO or K1328-KO cells, the proximal region of the DC was often located near the SDAs, suggesting that proteins present at these two locations may facilitate centriole cohesion in the absence of the CCDC61/K1328/CNTLN complex (Figure 7C). The position where NIN and CEP170 are detected corresponds approximately to the location of the central transverse striation of the BB-DC linker seen by electron microscopy. The antibodies we used target the C-terminal region of both proteins, which also allow them to interact with each other (Lamla, 2008). Consistent with this, a simultaneous labeling revealed that these two regions colocalize at the BB-DC linker (Supplemental Figure S5). NIN is a large protein (240 kDa) composed mainly of coiled-coil domains, which could adopt an elongated conformation. We thus compared the previous staining pattern with that obtained with an antibody directed against the N-terminal part of NIN. The two patterns were similar, however, indicating that the N-terminal and C-terminal domains both localize in the central region of the BB-DC linker (Supplemental Figure S5). To determine whether this might correspond to the central striation of the BB-DC linker, we first determined whether the striations could be visualized using pan-labeling with dye-conjugated NHS-ester (M’Saad & Bewersdorf, 2020). Transverse striations resembling those of the BB-DC linker were visible between the BB and DC in ciliated RPE1 cells, with a prominent striation located centrally as seen by electron microscopy (Figure 7D). We then co-localized CEP170 with the BB-DC linker. For this, we stained CEP170 after the pan-labeling to avoid revealing the antibodies with NHS-ester, as this would complicate the observation of the striations. We focused on CEP170 because the antigen recognized by our antibody contains few free primary amines and is therefore less likely to be modified by NHS-ester binding. Although pan-labelling was less effective in these conditions, we observed CEP170 localizing near the central striation of the BB-DC linker (Figure 7E). Together, these results suggest a model in which the positioning of the DC near the BB in ciliated cells involves the formation of the BB-DC linker, which contains the NIN and CEP170 proteins and is anchored at the DC by the CCDC61/K1328/CNTLN complex (Figure 7F).

**Figure 7:**
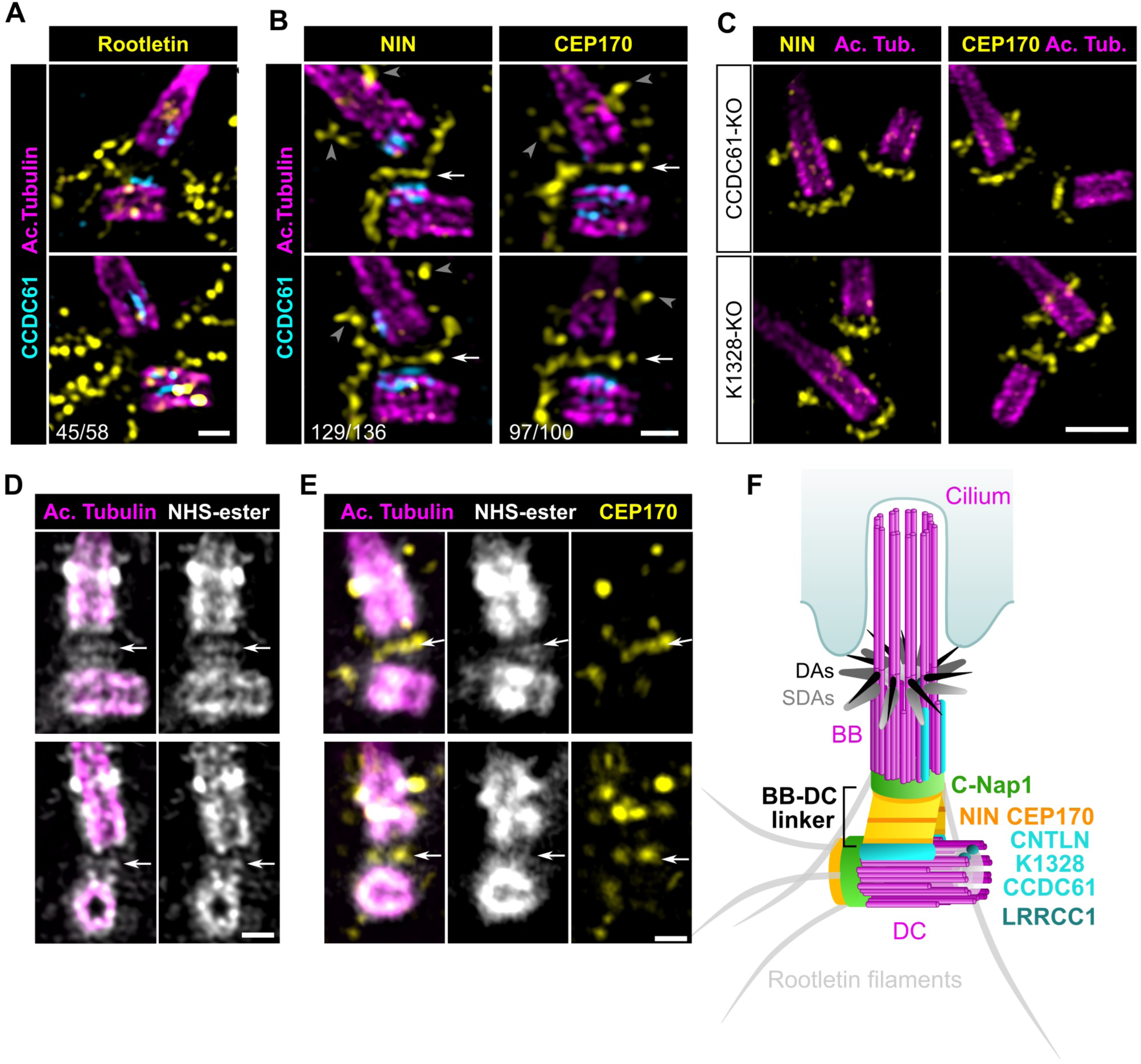
NIN and CEP170 are components of the BB-DC linker. **A)** U-ExM views showing the localization of rootletin (yellow) in WT ciliated RPE1 cells. CCDC61 is in cyan and acetylated tubulin in magenta. Scale bar, 200 nm. **B)** U-ExM views showing the localization of NIN or CEP170 (yellow) in WT ciliated RPE1 cells. The arrows show the linear pattern formed by NIN and CEP170 along the DC triplets labelled by CCDC61 (cyan). The arrowheads show SDAs. Acetylated tubulin is in magenta. Bar, 200 nm. **C)** U-ExM views showing NIN or CEP170 (yellow) in CCDC61-KO or K1328-KO ciliated cells. Even when the centrioles are nearby, the position of the DC at the proximal end of the BB and the localization of NIN or CEP170 along the side of the DC is lost. Acetylated tubulin is shown in magenta. Bar, 500 nm. **D)** U-ExM views of centrioles in WT ciliated RPE1 cells labelled with anti-acetylated tubulin antibody (magenta) then NHS-ester (gray). Arrows indicate the central striation of the BB-DC linker. Bar, 200 nm. **E)** U-ExM views of centrioles in WT ciliated RPE1 cells sequentially labeled with anti-acetylated tubulin antibody (magenta), NHS ester (gray), and anti-CEP170 antibody (yellow). Arrows indicate the central striation of the BB-DC linker. Bar, 200 nm. **F)** Diagram showing a model for the organization of the BB-DC linker in ciliated cells. The DC is connected laterally to the BB-DC linker via the CCDC61/K1328/CNTLN complex. NIN and CEP170 might form the two central layers of the linker. C-Nap1, known to recruit NIN and CEP170 to the proximal end of centrioles, likely anchors the BB-DC linker to the BB. The SDA pools of NIN and CEP170 are not highlighted.

## Discussion

Centriole rotational asymmetry is an ancient feature in eukaryotes that allows coordinating ciliary and cell polarity. Here, we outline the mechanisms underlying this asymmetry and its roles at the base of the primary cilium. We show that rotational asymmetry is established along centrioles through the coordinated recruitment of four proteins. LRRCC1 is recruited first and facilitates the recruitment of CCDC61 to two or three consecutive triplets at the distal end of procentrioles. In elongated centrioles, LRRCC1 remains in the distal lumen, whilst CCDC61 extends along the proximal half of the triplets, on their cytoplasmic side. CCDC61 then enables the recruitment of K1328 and CNTLN to the proximal region. The CCDC61/K1328/CNTLN complex allows the attachment of a linker connecting the proximal end of the BB to the lateral side of the DC in cells forming a primary cilium. The BB-DC linker allows the close positioning of the DC relative to the BB, a feature that is conserved in ciliated cells. Upstream of cilium formation, the CCDC61/K1328/CNTLN complex induces an asymmetry of the PCM, which likely facilitates the formation of the BB-DC linker during the following cell cycle. In MCCs, where BBs do not have an associated DC, the CCDC61/K1328/CNTLN complex shows the same localization pattern as in the centrosome, supporting a versatile role of this complex in anchoring asymmetric structures to centrioles and BBs.

### Conserved mechanisms govern the establishment of centriole rotational asymmetry

Despite the structural diversity of appendages that decorate centrioles asymmetrically in cells with motile cilia, the molecular mechanisms that determine the triplets on which these appendages are formed appear to be conserved. LRRCC1/VFL1 localizes to the distal end of the centrioles, opposite the direction of ciliary beating, in mouse MCCs and in *Chlamydomonas reinhardtii* (Silflow *et al*, 2001; Gaudin *et al*, 2022). Likewise, CCDC61/VFL3 localizes to the proximal part of the same triplets in MCCs, and is present opposite the direction of cilia beating in *Paramecium tetraurelia* (Bengueddach *et al*, 2017). We found that the respective localizations of LRRCC1 and CCDC61 are also conserved in the centrioles of the centrosome. This indicates that the mechanisms coordinating the recruitment of the different effectors of centriole rotational asymmetry are conserved throughout evolution and across all cell types. Our findings in cells forming a primary cilium are therefore also likely relevant for the polarization of motile cilia, with possible implications for the study of motile ciliopathies such as primary ciliary dyskinesia.

The first identified step in establishing centriole rotational asymmetry is the recruitment of LRRCC1 when procentrioles are on average ∼ 150 nm in length. This occurs after completion of the bloom phase, which involves the assembly of the cartwheel, MT triplets and the distal cap, and coincides with the onset of centriole elongation (Laporte *et al*, 2024). In *Chlamydomonas*, Vfl1p is present at the distal end of proBBs, which are arrested after the assembly of MT triplets and distal structures but prior to the elongation stage (Silflow *et al*, 2001; Geimer & Melkonian, 2005). The timing of LRRCC1 recruitment is thus overall conserved. CCDC61 is incorporated shortly after LRRCC1 at the distal end of elongating procentrioles and initially colocalizes with LRRCC1. CCDC61 can interact with LRRCC1, suggesting that LRRCC1 helps recruiting CCDC61 via a direct interaction between the two proteins. This is also supported by the observation that partial depletion of LRRCC1 decreases the amount of CCDC61 at the centrioles. As procentriole elongation takes place, LRRCC1 remains near the distal cap, while CCDC61 spreads along the proximal region. CCDC61, a paralog of SAS-6, can oligomerize to form linear filaments. A mutant version of *Chlamydomonas* Vfl3p that cannot oligomerize only partially rescues the *vfl3* mutant phenotype, but is correctly localized, suggesting that oligomerization is not essential for Vfl3p recruitment to the centrioles. In contrast, a mutation that prevents MT binding localizes abnormally and fails to rescue the *vfl3* mutant (Ochi *et al*, 2020). CCDC61 can interact with MTs (Ochi *et al*, 2020; Pizon *et al*, 2020), and our U-ExM data support that it is located close to centriole triplets. It is thus plausible that CCDC61 interacts directly with triplet MTs. Nevertheless, interactions with additional centriole components may explain why CCDC61 is confined to the proximal region. CCDC61 distribution along the proximal-distal axis is similar to that of the torus component CEP152. Components of the AC-linker, a protein complex that connects the A-tubule of a triplet to the C-tubule of its neighbor, are also similarly localized. The AC-linker stabilizes the centriole and is required for the recruitment of the torus, suggesting that the torus is anchored at the AC-linker (Laporte *et al*, 2024; Bournonville *et al*, 2025). Our observation that the CCDC61/K1328/CNTLN complex induces an asymmetry in the distribution of CEP152 and PCNT suggests that there may be competition between these proteins for binding to the centrioles. The recruitment of CCDC61/K1328/CNTLN on the one hand, and of CEP57/CEP63/CEP152 and CEP57/PCNT on the other (Watanabe *et al*, 2019; Ito *et al*, 2021; Zhao *et al*, 2020; Wei *et al*, 2020), may involve common partners, possibly within the AC-linker. CCDC61 is essential to the recruitment of K1328, and K1328 is required for recruiting CNTLN. CNTLN contains a MT binding domain located in the C-terminal region, which might help anchor the complex to the triplets (Jing *et al*, 2016). Interestingly, the antibody we used, which recognizes the N-terminus of CNTLN, consistently produced a pattern similar to that of CCDC61, but shifted clockwise by approximately one triplet (in distal view). This suggests that the CCDC61/K1328/CNTLN complex could either bridge adjacent triplets or bind at different sites along the AC linker. The presence of K1328 and CNTLN in the BBs of MCCs indicates that the complex’s function is conserved at the base of motile cilia. Orthologs of K1328 are also present in protist genomes including *Chlamydomonas* and the ciliate *Tetrahymena thermophila* (Supplemental Figure S3). We could not identify clear orthologs of CNTLN in protists, but this was not unexpected given that this protein shows little amino acid conservation, even among animals. Thus, the whole CCDC61/K1328/CNTLN complex is possibly ancestral in eucaryotes.

### The BB-DC linker enables DC repositioning near the base of the primary cilium

Data from the literature and our own observations show that the BB-DC linker is widespread in mammalian tissues and has a consistent architecture (Yamamoto & Kataoka, 1986; Hagiwara *et al*, 2002, 2008; Gallagher, 1980; Webber & Lee, 1975; Wheatley, 1967). The BB-DC linker differs from the rootletin-containing centrosome linker in terms of structure and molecular composition, but it shares similarities with SDAs. CEP170 and NIN are components of both SDAs and the BB-DC linker. In SDAs, CEP170 and NIN are located at the periphery of a filamentous stem (LeGuennec *et al*, 2021; Nguyen *et al*, 2020; Chong *et al*, 2020; Carden *et al*, 2023). NIN adopts a semi-elongated conformation, with its N-terminal and C-terminal ends located at the periphery of SDAs, and the central portion further inward, with approximately 60 nm separating the N-terminus from the central region (Nguyen *et al*, 2020). NIN, which consists primarily of coiled-coil domains, may thus fold back to form filaments. In the BB-DC linker, CEP170 and both ends of NIN are located near the central striation of the BB-DC linker, between two layers of 60 nm-long filaments that resemble the ones found in SDAs (Carden *et al*, 2023). The two layers appear similar and may therefore have the same composition. In that case, they would be in opposite orientation, with CEP170 and the terminal domains of NIN near the central striation, and the central domain of NIN in the filamentous portions on either side. On the BB side, the filamentous layer is separated by a transverse striation from a less structured layer located immediately at the proximal end of the BB. This layer likely contains C-Nap1, which recruits NIN and CEP170 to the proximal end of the centrioles (Mazo *et al*, 2016). Proximal NIN was shown to play a role in centriole cohesion in RPE1 cells, and it may contribute to this process in different ways during the various phases of the cell cycle (Theile *et al*, 2023). Nevertheless, intercentriolar distance in NIN-depleted ciliated cells is similar to that in CCDC61-KO, K1328-KO and CNTLN-KD, which supports that NIN acts primarily through the BB-DC linker to reposition the DC close to the BB in this context. NIN and CEP170 are present at the proximal ends of centrioles throughout interphase, whereas the BB-DC linker is only observed in ciliated cells. This suggests that proximal proteins may be rearranged or change conformation during formation of the BB-DC linker. Future work will also need to examine the possible involvement of proteins with a similar distribution to NIN and CEP170—such as Kif2A, p150^glued^, CCDC120, and CCDC68—in the formation of the BB-DC linker (Mazo *et al*, 2016; Huang *et al*, 2017).

On the DC side, the BB-DC linker is anchored by the CCDC61/K1328/CNTLN complex. CNTLN was previously involved in centriole cohesion, but this was attributed to a role in forming the centrosome linker. This was supported by the fact that CNTLN interacts with CEP68, a component of rootletin filaments (Fang *et al*, 2014; Vlijm *et al*, 2018). We did not detect CNTLN at the proximal tip of centrioles or in between centrioles, but is possible that a small fraction is associated to rootletin filaments and was not detected by U-ExM. An intriguing question is why the linker forms only between the proximal end of the BB and the DC side, even though the proteins involved in its assembly are present on both centrioles. One hypothesis is that some of these proteins might undergo changes in their levels or post-translational modifications. Consistent with this idea, CCDC61, K1328, and CNTLN are, on average, twice as abundant at the DC as at the BB in ciliated cells, which could contribute to the asymmetry in linker formation.

The positioning of the DC close and orthogonal to the proximal end of the BB is conserved in evolution, already occurring in early-branching animals and in choanoflagellates, the closest unicellular relative of animals (Pozdnyakov *et al*, 2021; Karpov, 2016). This configuration is also observed in mammalian sperm, where it plays a key role in the formation of a structure that connects the flagellum to the nucleus, known as the head-to-tail coupling apparatus (Takeda *et al*, 2025). Interestingly, CNTLN is a component of the head-to-tail coupling apparatus, and its inactivation in mice leads to sperm decapitation and male infertility (Zhang *et al*, 2021). However, it is not clear at this stage whether its precise localization corresponds to what we observed at the base of the primary cilium. In RPE1 cells, the proximity of the DC to the BB promotes the formation of the primary cilium by allowing the localization of the ubiquitin ligase cofactor Neurl4 at the BB (Loukil *et al*, 2017). Defects in primary cilium formation have been reported for CCDC61-KO and CNTLN-KO RPE1 cells in some instances – although not in the CRIPSPR clones used in the present study (Pizon *et al*, 2020) (Supplemental Figure S3), suggesting that ciliary assembly is not optimal in the absence of the BB-DC linker (Ochi *et al*, 2020; Li *et al*, 2023). DC repositioning could affect the organization of the cytoskeleton or rootletin filaments, which were shown to contribute to ciliogenesis by recruiting centriolar satellites (Turn *et al*, 2021). Other aspects, such as cilium stability or ciliary signaling, could also depend on the organization of the ciliary base (Chen *et al*, 2015). Future work will shed light on the role of DC repositioning in ciliated cells and provide insight into its evolutionary conservation.

## Methods

### Cell culture

RPE1 cells (hTERT-RPE1, RRID:CVCL_4388; American Type Culture Collection, authenticated by STP profiling, no mycoplasma detected) were cultured in DMEM/F-12 medium (ThermoFisher Scientific) supplemented with 10 % fetal calf serum (ThermoFisher Scientific), 100 U/mL penicillin and 100 μg/mL streptomycin (ThermoFisher Scientific). Ciliogenesis was induced by culturing RPE1 cells in medium without serum during 48 hours. U2-OS cells were cultured in DMEM medium (ThermoFisher Scientific) supplemented with 10 % fetal calf serum and antibiotics as previously. All cells were kept at 37°C in the presence of 5 % CO_2_.

### Mouse tissues

All experiments were performed in accordance with French Agricultural Ministry and European guidelines for the care and use of laboratory animals. Salivary glands, spleen and trachea fragments were isolated from wild-type mice of the Swiss background—one male and two females.

### CRISPR/Cas9 editing

The CCDC61-KO RPE1 clone (clone 28.1) and the LRRCC1 partially depleted clone (clone 1.1) used in this study were described previously (Pizon *et al*, 2020; Gaudin *et al*, 2022). The K1328-KO and CNTLN-KD clones were obtained using the plasmid pSpCas9(BB)-2A-GFP (PX458), a gift from Feng Zhang (Addgene plasmid #48138; http://n2t.net/addgene:48138; RRID:Addgene_48138). RPE1 cells were transfected with a mix of plasmids containing the following sgRNAs: 5’- TGG GCA TCT CTG GTG CAT GG - 3’ (sg1) and 5’- CTA GGA GTA GGC TTA AGA TG - 3’ (sg2) for K1328; 5’- AGT ACA CGC AAT GCG CAG CG - 3’ (sg1), 5’- AGA GCT GAG CAG CCT AAA GG - 3’ (sg2) and 5’- AAG TGA CCA AAA CCA AAC CC - 3’ (sg3) for CNTLN. Two days after transfection, individual GFP-positive cells were sorted by flow cytometry in 96-well plates. Clones were picked after 2 weeks and analyzed by PCR to detect insertions/deletions. The mutant clones were further characterized by genomic sequencing as well as transcript sequencing in the case of CNTLN-KD clones. Genomic DNA was isolated using Quick Extract DNA extraction solution (Biosearch Technologies). For transcript analysis, total RNA was isolated using the Nucleospin RNA kit (Macherey-Nagel) and cDNAs were obtained using oligo-d(T) and Superscript III reverse transcriptase (Thermofisher Scientific) following recommendations of the manufacturers.

### RNAi

Ready to use double-stranded siRNAs were used to further reduce LRRCC1 levels in the hypomorph CRISPR clone 1.1 (target sequence: 5’- AAG GAG AAA GAT GGA GAC GAT - 3’) and to deplete NIN (target sequence: 5’- AAG AAG AAC TGG AAC GTT GTA - 3’) and were purchased from Qiagen. siRNA delivery was performed using Lipofectamine RNAiMAX diluted in OptiMEM medium (ThermoFisher Scientific). For LRRCC1 depletion, cells were processed for U-ExM after 48 hours. For NIN depletion, cells were switched to serum-free medium after 48 hours to induce ciliogenesis. After 24 hours in serum-free medium, cells were fixed in methanol at -20°C during 5 minutes and processed for immunofluorescence.

### MT targeting assay

The GFP-CCDC61 and LRRCC1-GFP constructs, cloned into pEGFP-C2 (Clontech) and pCDNA-5FRT (ThermoFisher Scientific) vectors, respectively, were described previously (Gaudin *et al*, 2022; Pizon *et al*, 2020). The LRRCC1-GFP construct corresponds to full-length LRRCC1 (NP_208325.3) with GFP inserted at after aa 402. The LRRCC1^Δ900-930^-GFP construct was obtained by deleting the CCDC61-interaction domain predicted by AlphaFold3 from the LRRCC1-GFP construct. LRRCC1 fragments aa 1-205, 195-415, 405-1032 and 405-1032Δ900-93 (Supplemental Figure S2) were cloned into the peGFP-C2 vector downstream of GFP. The full-length coding sequence of K1328 (NP_065827) or fragments 1-289 and 290-577 were cloned into the pcDNA5/FRT/TO-neo vector (1795), a gift from Jonathon Pines (Addgene plasmid # 41000; http://n2t.net/addgene:41000; RRID:Addgene_41000). Fragments corresponding to full-length CNTLN (NP_060208.2) or protein domains were cloned into the pEGFP-C2 vector. To target CCDC61 to MTs, the coding sequence for the full-length protein or protein domains were inserted upstream of mCherry and the MT-binding domain of FAM161A (MBD, amino acids 176–479 of NP_001188472.1) into a vector derived from pEGFP-C2 in which EGFP was replaced by mCherry. For targeting CNTLN to the MTs, the MBD sequence was inserted upstream of mCherry and CNTLN sequences in the mCherry vector derived from pEGFP-C2. The constructs were transfected into U2-OS cells grown on glass coverslips using Lipofectamine 2000 (ThermoFisher Scientific) following the manufacturer’s recommendations. 24h after transfection, cells were fixed with 4% paraformaldehyde in PBS during 10 min at RT and processed for immunofluorescence.

### Antibodies

Fragments encoding either aa 961-1032 of LRRCC1 (NP_208325.3), aa 337-512 of CCDC61 (NP_001254652) or aa 91-200 of K1328 (NP_065827) were cloned in pGST-Parallel1 and expressed in *Escherichia coli.* The GST-fusion proteins were purified under native conditions using glutathione agarose (ThermoFisher Scientific) and the LRRCC1 fragments were recovered by Tev protease cleavage and dialyzed before rabbit (K1328) or goat (LRRCC1 and CCDC61) immunization (Covalab). Antibodies were affinity-purified over the corresponding GST fusion proteins bound to Affi-Gel 10 resin (Bio-Rad). Other primary and secondary antibodies used in this study are listed in Table 1.

**Table 1.**
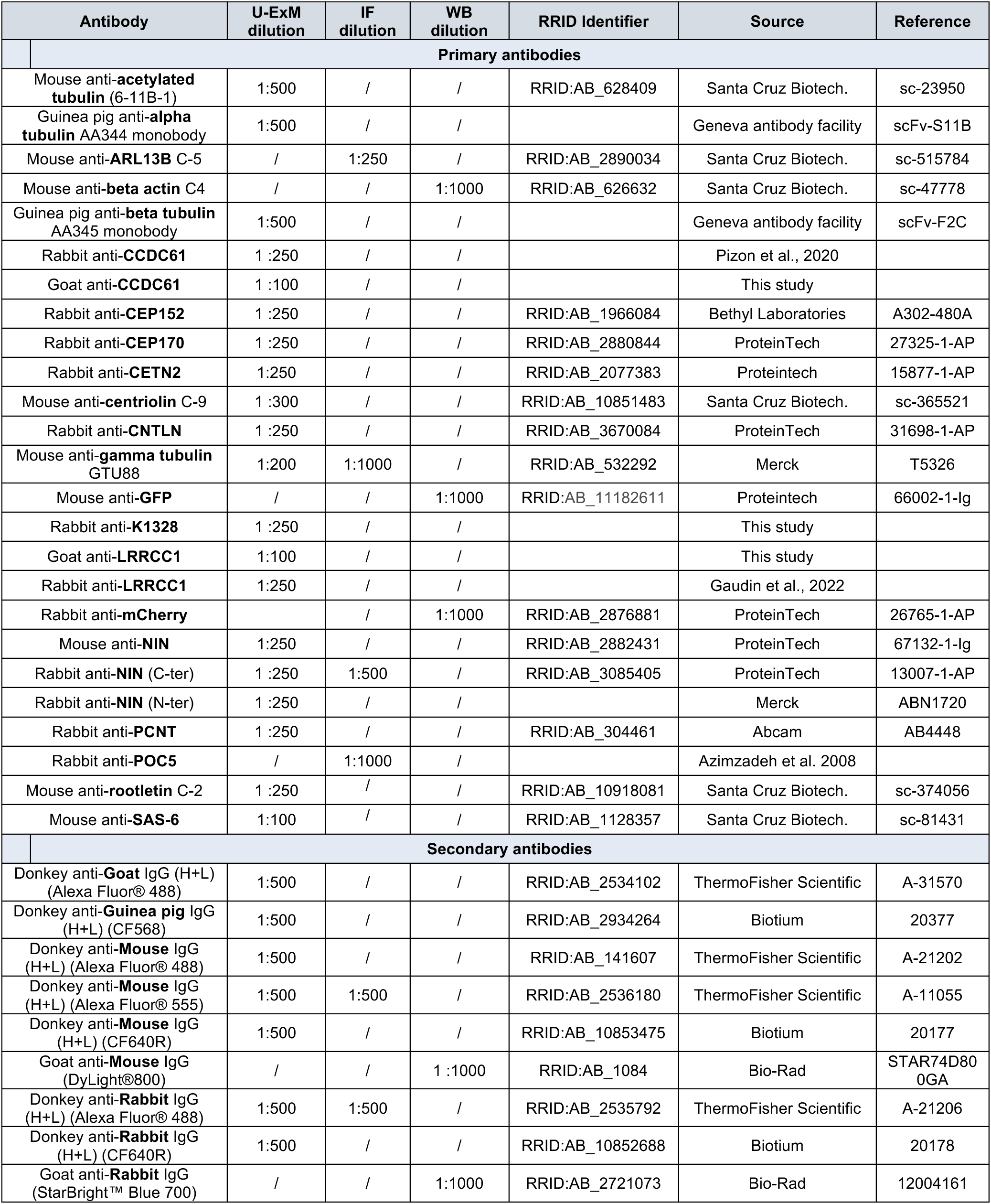
Antibodies used in this study.

### Immunofluorescence

To analyze intercentriolar distance and ciliogenesis efficiency, cells at ∼ 80% confluence were cultured for 24 hours in serum-free medium to induce ciliogenesis, then fixed in methanol at -20°C for 5 minutes. Non-specific sites were blocked for 10 minutes in PBS containing 0.01% Tween-20 (PBST) and 3% BSA, then incubated for 1 hour with anti-POC5 and anti-ARL13B antibodies diluted in PBST-BSA to label centrioles and cilia, respectively. After washing 3 x 1 minute in PBST, cells were incubated 1 hour with secondary antibodies in PBST-BSA containing 5 µg/mL Hoechst 33342 (ThermoFischer Scientific), washed in PBST as previously, and mounted using Fluoromount-G (ThermoFischer Scientific). Images were acquired using an Axio Observer Z.1 microscope (Zeiss) equipped with a sCMOS Orca Flash4 LT camera (Hamamatsu) and a 63x objective (Plan Apo, N.A. 1.4).

### Ultrastructure expansion microscopy

We used the U-ExM protocol described in (Gambarotto *et al*, 2019) with slight modifications. Cells grown on glass coverslips were incubated in a fresh solution of 1 % acrylamide and 0.7 % formaldehyde diluted in PBS. After incubating overnight at 37 °C, the coverslips were washed with PBS and placed cells down on a drop of 35 μL monomer solution (19.3 % sodium acrylate, 10 % acrylamide, 0.1 % bis-acrylamide in PBS) to which 0.5 % TEMED and 0.1 % ammonium persulfate were added just before use. The coverslips were incubated 5 minutes on ice then 1 hour at 37°C, then transferred to denaturation buffer (200 mM SDS, 200 mM NaCl, 50 mM Tris pH9) for 15 minutes with agitation to detach the gels from the coverslips. The gels were then incubated in denaturation buffer 1.5 hours at 95 °C, washed 2 x 30 minutes in deionized water then incubated overnight in water at room temperature to allow expansion of the gel. The gels were measured at this step to determine the coefficient of expansion. After 2 x 10 minutes in PBS, the gels were cut into smaller pieces then incubated 3 hours at 37 °C with primary antibodies diluted in saturation buffer (3 % BSA, 0.05 % Tween-20 in PBS). The gel fragments were then washed 3 x 10 minutes in PBST-0.1%, incubated 3 h with secondary antibodies and washed in PBST-0.1% as previously. For NHS-ester staining, gels were incubated 1.5 hours with 100 µg/mL CF®640R Dye SE/TFP Ester (Biotium) diluted in PBS, then washed 4 x 15 minutes in PBST-0.1%. Finally, the gels were incubated 2 x 30 minutes in de-ionized water then left to expand overnight in de-ionized water to regain their maximum size. For U-ExM of mouse trachea, spleen and salivary glands, tissue explants were fixed with 4% paraformaldehyde in PBS before being processed for U-ExM. Gels were imaged on Lab-Tek chamber slides (0.15 mm) coated with poly-lysine (ThermoFisher Scientific). Images were acquired at room temperature using an LSM980 confocal microscope with Airyscan 2 (Zeiss) equipped with an oil 63x objective (Plan Apo, N.A. 1.4).

### Image analysis

Protein levels were determined using ImageJ software (Schneider *et al*, 2012) by measuring the fluorescence intensity at individual centrioles and subtracting the cytoplasmic background. Images of individual centrioles in U-ExM are maximum intensity projections of all z-sections comprising the signal of interest (z-increment ∼ 0.15 µm). Average images were obtained as described previously (Gaudin *et al*, 2022). Briefly, stacks of images were acquired for centrioles nearly perpendicular to the imaging plane to maximize the resolution in transverse views. Calculating the average image consisted of several steps: cropping out individual centrioles, aligning them, providing reference points, standardizing centrioles using the reference points, and averaging, as described previously. For alignment of tracheal cell centrioles, since the graphical user interface can only accommodate two channels, the position of the basal foot provided by the γ-tubulin channel was reported manually in the acetylated tubulin channel using Image J. The images were then processed as before using the manual annotation as a reference point for the basal foot.

### Immunoprecipitation and Western blot

HEK293 cells were transfected with plasmids encoding CCDC61, K1328 and CNTLN fused to GFP or mCherry (see MT targeting assay) using PEI (Sigma) as described previously. 24h after transfection, cells were resuspended in lysis buffer (50 mM Tris pH 7.5, 150 mM NaCl, 1 % Igepal-CA-630, 1 % sodium deoxycholate, 0.1 % SDS, 0.5 mM EDTA, 1 mM MgCl_2_) supplemented with 80 U/mL DNAse I (Roche), PhoSTOP phosphatase inhibitor (Roche) and cOmplete protease inhibitor cocktail (Roche). After 30 minutes on ice, the lysates were centrifuged at 17 000 g for 10 minutes at 4°C. The supernatants were diluted 1.5 x with dilution buffer (50 mM Tris pH 7.5, 150 mM NaCl, 0.5 mM EDTA) then incubated with GFP-Trap magnetic agarose beads (ChromoTek) and rotated for 1.5 hours at 4°C. After washing 4 x with wash buffer (50 mM Tris pH 7.5, 150 mM NaCl, 0.05 % Igepal-CA-630, 0.5 mM EDTA), immunoprecipitated proteins were recovered by resuspending the beads in Laemmli buffer and heating at 95 °C for 10 minutes. The samples were then run on 4-20% Mini-Protean TGX precast protein gels (Bio-Rad) and transferred onto PVDF membrane using the iBlot 2 blot system (ThermoFischer Scientific). The membranes were blocked and incubated with mouse anti-GFP and rabbit anti-mCherry antibodies then fluorescent secondary antibodies, and visualized ChemiDoc MP imaging system (Bio-Rad).

### Centrosome purification

Centrosomes were purified from serum starved RPE1 cells using CAPture, as described previously (Carden *et al*, 2023). Briefly, RPE1 cells, both wild type and CCDC61-KO line (Ochi *et al*, 2020), were grown on 4x 15cm dishes in DMEM F-12 GlutaMAX (Thermo Fisher Scientific) supplemented with 10% FBS till reaching confluence and ciliated by further incubating them in the growth media without FBS for 48 hours. The cells were collected using a scraper in PBS and pelleted using centrifugation before being stored at -80 ℃. 50 µl Dynabeads M-280 Streptavidin (Thermo Fisher Scientific) per pulldown were washed three times in TBSN (TBS with 0.1 %(v/v) NP-40) and suspended in 100 µl of TBSN. 5 µl of 5 mg/ml CAPture peptide suspended in TBS were added to the washed beads and incubated on a rotating platform at 12 rpm for 60 min at 4 ℃. While incubating, a cell pellet of the RPE1 cells were suspended in 1.5 ml of CAPture buffer (50 mM Tris-HCl pH 8.0, 300 mM NaCl, 5 mM NaF, 0.2 %(v/v) NP-40, 10 %(v/v) glycerol, complete EDTA-free protease inhibitor cocktail (Roche Diagnostics) on ice and sonicated briefly (2 x 3 second pulses) with an amplitude of 30% using a 3mm microtip probe, and centrifuged at 1800 g for 10 min at 4 ℃. The beads were then washed three times with TBSN followed once by CAPture buffer. The beads were added to the supernatant of the RPE1 lysate and incubated on a rotating platform at 12 rpm for 2 hours at 4 ℃. The beads were washed five times with CAPture buffer and transferred to a new 1.5 ml tube. 30 µl of the CAPture elution peptide (3xFLAG-hCCDC61^334-366^ (Carden *et al*, 2023) were added to the beads and left for 30 min at room temperature. The supernatant was collected and used for mass spectrometry or electron microscopy.

### Mass spectrometry

Gel samples were destained with 50% v/v acetonitrile and 50 mM ammonium bicarbonate, reduced with 10 mM DTT, and alkylated with 55 mM iodoacetamide. Digestion was with 6 ng/μl trypsin (Promega, UK) overnight at 37°C, and peptides extracted in 2% v/v formic acid 2% v/v acetonitrile, and analysed by nano-scale capillary LC-MS/MS using an Ultimate U3000 HPLC (ThermoScientific Dionex, San Jose, USA) to deliver a flow of approximately 300 nL/min. A µ-precolumn cartridge C18 Acclaim PepMap 100 (5 µm, 300 µm x 5mm (ThermoScientific Dionex, San Jose, USA), trapped the peptides prior to separation on a C18 Acclaim PepMap100 3 µm, 75 µm x 250 mm (ThermoScientific Dionex, San Jose, USA). Peptides were eluted with a 65-minute gradient of acetonitrile (5 to 40%). The analytical column outlet was directly interfaced via a modified nano-flow electrospray ionisation source, with a hybrid linear quadrupole ion trap mass spectrometer (Orbitrap QExactive, ThermoScientific, San Jose, USA). Data dependent analysis was carried out, using a resolution of 60,000 for the full MS spectrum, followed by ten MS/MS spectra in the linear ion trap. MS spectra were collected over a m/z range of 200–1800. MS/MS scans were collected using threshold energy of 35 for collision induced dissociation. LC-MS/MS data were then searched against the uniprot database, using the Mascot search engine programme (Matrix Science, UK). Database search parameters were set with a precursor tolerance of 10 ppm and a fragment ion mass tolerance of 0.8 Da. One missed enzyme cleavage was allowed and variable modifications for oxidized methionine, carbamidomethyl and phosphorylated serine, threonine and tyrosine were included.

### Cryo-electron tomography

The tomograms of the BB-DC pairs purified from RPE1 cells were collected as described previously (Carden *et al*, 2023). 6 µl of 10 nm gold (BBI Solutions) were added to 30 µl of the eluted basal bodies. 3,5 µl of the sample were blotted to a 200-mesh holey carbon R2/1 copper grid (Quantifoil) using a Vitrobot (Thermo Fisher Scientific). We used a FEI Titan Krios 2 microscope inserted the Volta phase plate (Danev & Baumeister, 2016) and equipped with a Gatan K2 Summit direct electron detector with energy filter (20 eV slit width) at MRC LMB. Tilt series were collected using SerialEM (Mastronarde, 2005) at 33,000x nominal magnification (3.7 Å / pixel) on the counting mode and at 3° increments with the dose-symmetric scheme between -60° and 60° tilt angles. The defocus was set at -0.75 µm. 10 movie frames per tilt angle were collected with dose of 0.25 e^-^/Å/frame. Alignframes in Etomo (Kremer *et al*, 1996) was used to align movie frames. Tomogram reconstruction was performed using Etomo from IMOD using weighted back-projection. Contrast transfer function estimation and correction were performed using NovaCTF (Turoňová *et al*, 2017). The reconstructed tomograms were binned by a factor of 4 and applied a Gaussian filter using Chimera (Pettersen *et al*, 2004). The contrast of the filtered tomograms weas further enhanced by averaging ten tomogram slides using the newstack module of IMOD.

### Statistical analysis

All statistical analyses were performed using the Prism 10 for Mac OS X software (GraphPad Software, Inc.). The number of experimental replicates and the statistical test used are indicated in the figure legends, and the p values are included when statistically different.

## Data availability

Data have not been deposited in external repositories at this stage.

## Author contributions

**Meriem Boumendjel**: Formal analysis; Funding acquisition; Investigation. **Griselda Wentzinger**: Formal analysis; Investigation. **Mohamed Bahida**: Formal analysis; Investigation. **Tamara Advedissian**: Formal analysis; Investigation. **Tania Joanet**: Formal analysis; Funding acquisition; Investigation. **Futura Gattobigio**: Investigation. **Farida Begum**: Formal analysis; Investigation. **Nicolas Moisan**: Formal analysis. **Mark van Breugel**: Conceptualization; Funding acquisition. **Takashi Ochi**: Conceptualization; Formal analysis; Funding acquisition; Investigation; Visualization. **Juliette Azimzadeh**: Conceptualization; Formal analysis; Funding acquisition; Investigation; Supervision; Visualization; Writing – Original Draft Preparation.

## Acknowledgements

We acknowledge the core imaging facility of Institut Jacques Monod (ImagoSeine facility, member of the France BioImaging infrastructure supported by grant ANR-10-INBS-04 from the French National Research Agency). We also thank Vincent Maupu-Massamba and Patricia Moussounda for their technical assistance. This work was supported by funding from Fondation ARC pour la recherche sur le cancer, Labex Who Am I? (supported by grants ANR-11-LABX-0071 and ANR-11-IDEX-0005-02) and ANR-21-CE13-008 and ANR-25-CE13-2931-01 to JA. M.v.B. was supported by the BBSRC (BB/X013030/1). MBo was a recipient of a MESRI PhD fellowship from the French government and a 4^th^ year PhD fellowship from La Ligue Contre le Cancer, and TJ from a full PhD fellowship from La Ligue Contre le Cancer.

## Abbreviations

BB: basal body
DC: daughter centriole
MC: mother centriole
MCC: multiciliated cell
MT: microtubule
SDA: subdistal appendage
U-ExM: ultrastructure expansion microscopy

