## Supplemental Material for "Rotational asymmetry is required to position centrioles at the base of the primary cilium"

### Supplemental Figure S1

**A**

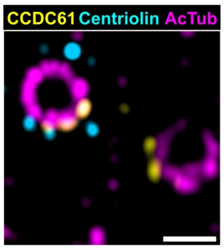


**B**

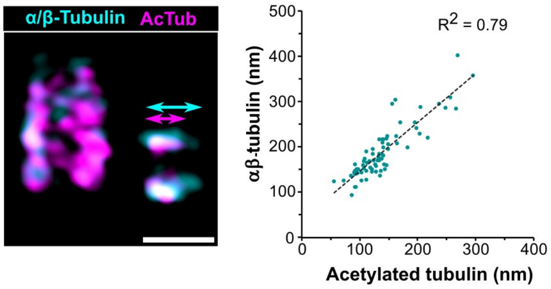

### Supplemental Figure S2

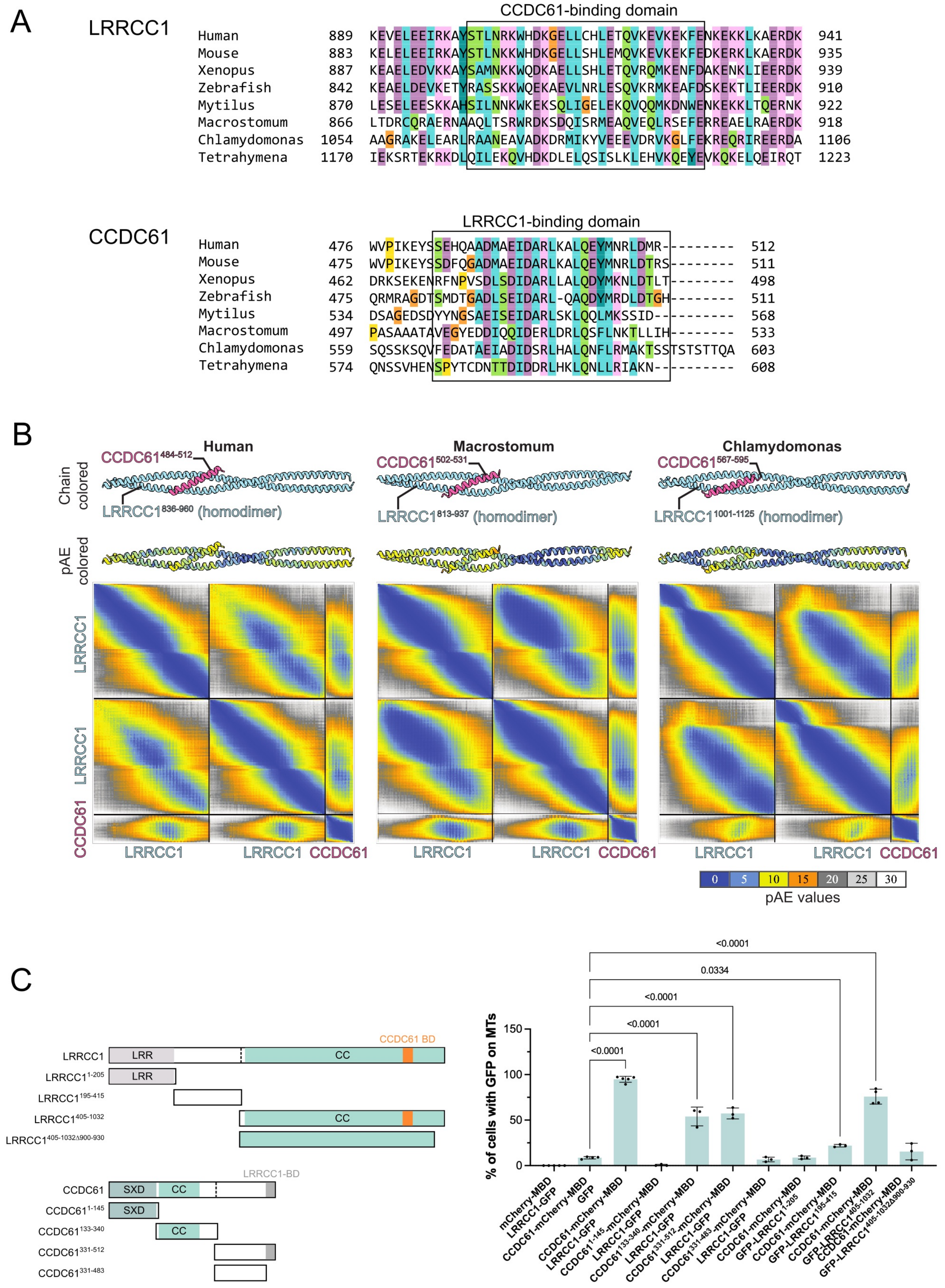

### Supplemental Figure S3

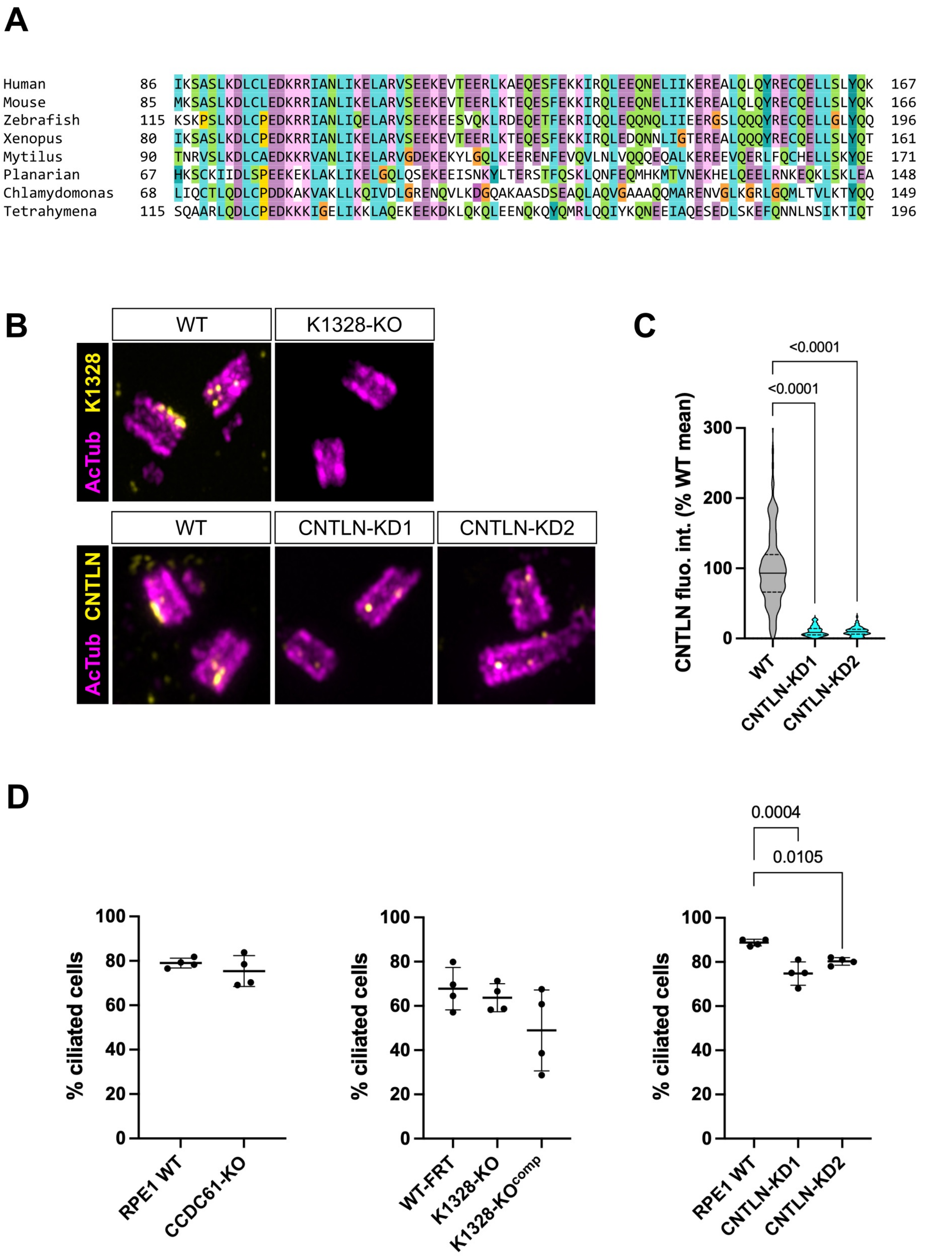

### Supplemental Figure S4

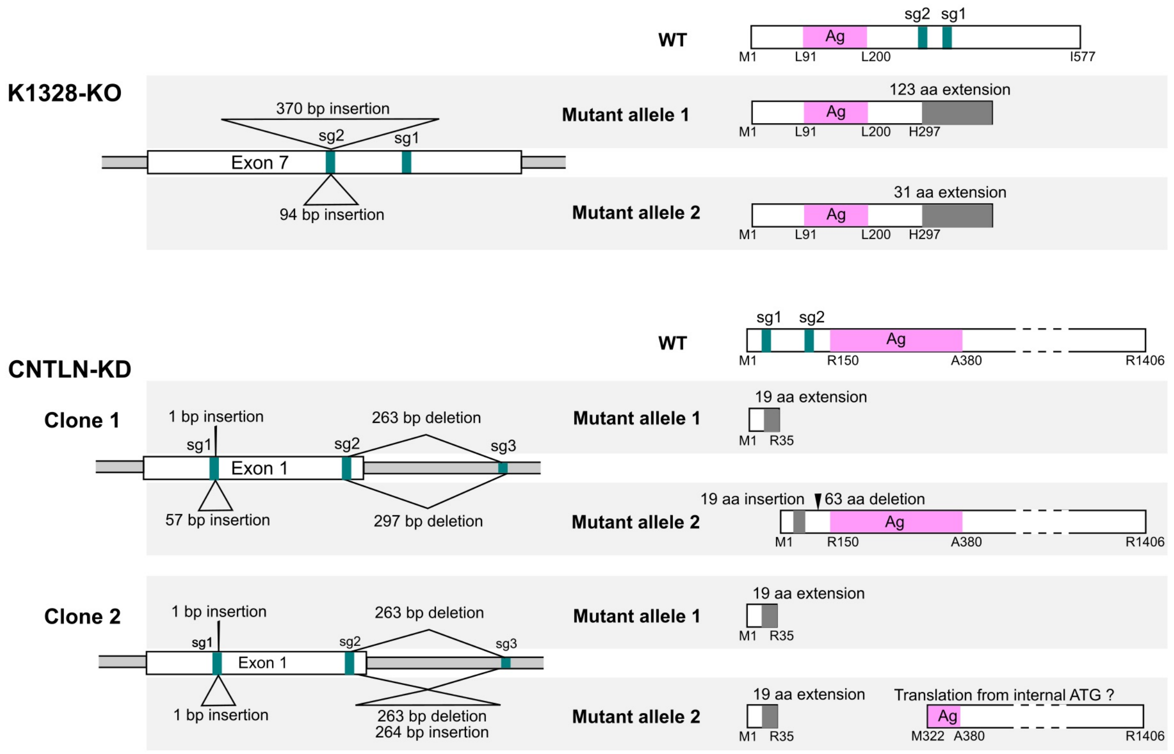

### Supplemental Figure S5

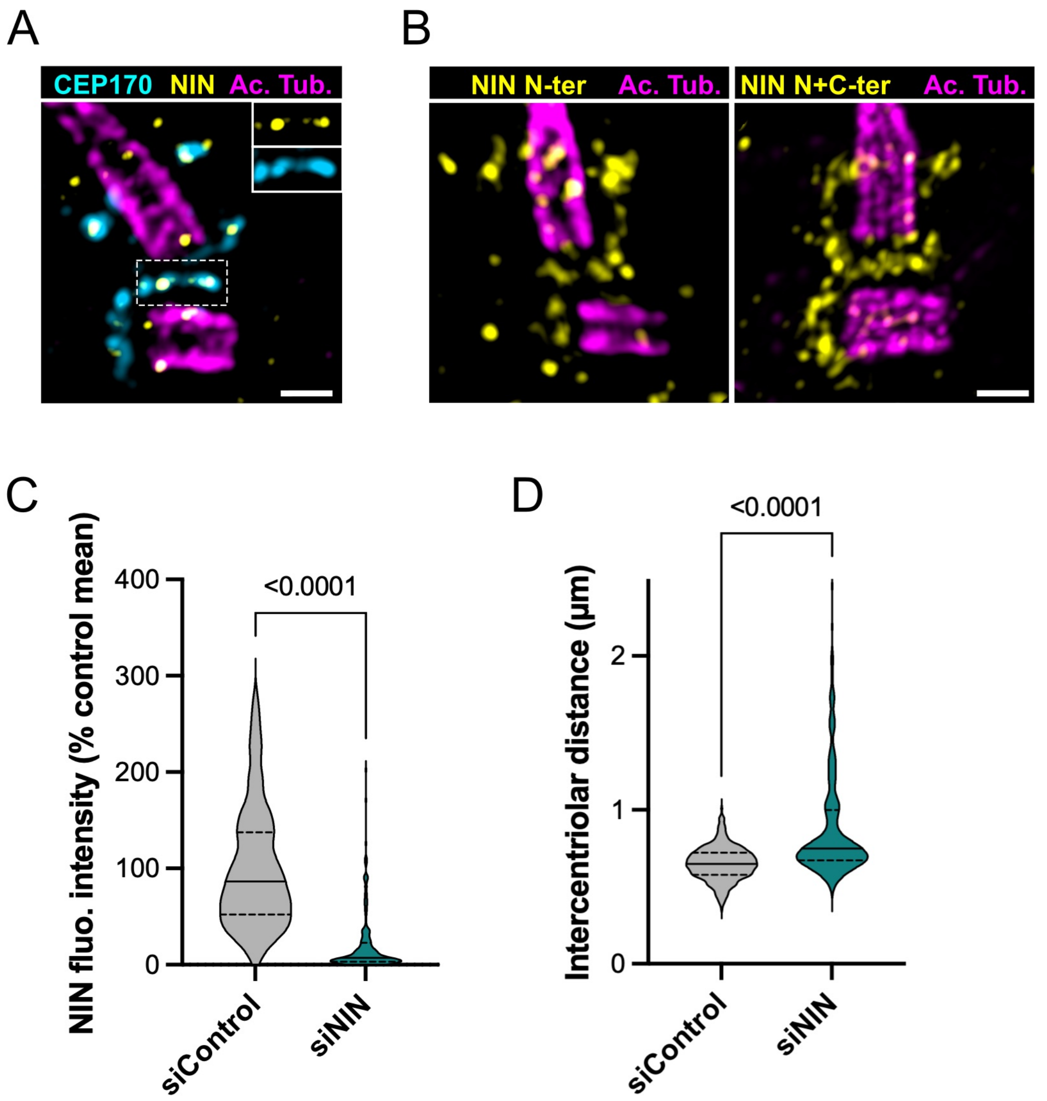

#### Supplemental Figure legends

**Supplemental Figure S1: A)** U-ExM views of RPE1 centrioles stained for CCDC61 (yellow), centriolin (cyan) and acetylated tubulin (magenta). Centriolin allows to distinguish MC and DC. Bar, 200 nm. **B)** Correlation between the length of acetylated tubulin staining (magenta) and the total triplet length determined by $\alpha/\beta$-tubulin staining (cyan) in RPE1 procentrioles. N = 75. Bar, 200 nm.

**Supplemental Figure S2: A)** Clustal Omega alignment of interaction domains between LRRCC1 and CCDC61. The following sequences were used: LRRCC1: human (Q9C099), mouse (Q69ZB0), *Xenopus laevis* (Q6NRC9), zebrafish (F1R597), *Mytilus coruscus* (A0A6J8A6D3), *Macrostomum lignano* (A0A267EDM9), *Chlamydomonas reinhardtii* (Q9SWH3), *Tetrahymena thermophila* (Q22WE5). CCDC61: human (Q9Y6R9), mouse (Q3UJV1), *Xenopus laevis* (A0A974C710), zebrafish (Q08CF3), *Mytilus edulis* (A0A8S3S6R3), *Macrostomum lignano* (A0A267FEW3), *Chlamydomonas reinhardtii* (Q68UI8), *Tetrahymena thermophila* (I7ME73). **B)** AlphaFold3 models of LRRCC1 homodimer in complex with CCDC61. The cartoon models were colored by protein and pAE values, and shown together with the pAE value heat maps of the models. The averaged ipTM values for the CCDC61-LRRCC1 homodimer in human, *Macrostomum*, and *Chlamydomonas* are 0.56, 0.58, and 0.52, respectively. **C)** MT targeting assay with subdomains of CCDC61 and LRRCC1 in U2-OS cells. The interaction domains predicted by Alphafold3 are highlighted. MBD: MT-binding domain of FAM161A used to target the constructs to MTs. CC: coiled-coil; SXD: domain shared with SAS-6 and XRCC4 (Ochi *et al*, 2020). LRR: leucine-rich repeat. Three independent experiments, N > 80 for each condition. Bars indicate standard deviation. p values are indicated when significantly different from the MBD-CCDC61-mCherry/GFP control (One-way ANOVA).

**Supplemental Figure S3: A)** Clustal Omega alignment of K1328 most conserved region. The following sequences were used: human (Q86T90), mouse (Q6NZK5), *Xenopus laevis* (A0A8J1M4I7), zebrafish (A4IG80), *Mytilus coruscus* (A0A6J8CGQ4), *Schmidtea mediterranea* (SMED30034649), *Chlamydomonas* sp. UWO 241 (KAG1671402.1), *Tetrahymena thermophila* (Q22DH6). **B)** U-ExM views of centrioles from WT, K1328-KO, or CNTLN-KD cells labeled with anti-K1328 or anti-CNTLN (yellow) and anti-acetylated tubulin (magenta) antibodies. K1328 is undetectable at the centrosome in K1328-KO cells, while residual CNTLN labeling is detected in both CNTLN-KD clones. **C)** Quantification of residual CNTLN signal in CNTLN-KD clones as a percentage of the mean control value. Three independent experiments, N > 98 for each condition. p values are indicated when significantly different from the control (One-way ANOVA). **D)** Percentage of ciliated cells after 24 hours in serum-free medium. Three independent experiments, N > 1000 for each condition. Bars represent the mean and standard deviation. p values are indicated when significantly different from the control (CCDC61: Two-tailed T-test; CNTLN, K1328: One-way ANOVA).

**Supplemental Figure S4:** Analysis of the genomic sequence of the CRISPR clones generated for this study. The K1328-KO clone carries insertions in both *K1328* alleles (top left), leading to a frameshift after H297. The corresponding truncated proteins (top right), which still contain the antigen used for generating our anti-K1328 antibody (in magenta), are not detected at the centrosome. The CNTLN-KD clones contain an insertion at the first CRISPR target site (sg1) and a deletion between the other two that deletes the end of exon 1 on both alleles (bottom left). Allele 2 of clone 2 also has an insertion between sg2 and sg3. A transcript corresponding to allele 1 of clone 1 is detected by RT-PCR, which theoretically allows the synthesis of a protein with an insertion of 19 amino acids and a deletion of 63 amino acids in its N-terminal region due to alternative splicing within exon 1 (bottom right). A transcript is also detected for allele 2 of clone 2, which should lead to a stop after R35. The presence of residual protein at the centrosome may be due to internal re-initiation of translation from M322.

**Supplemental Figure S5**: **A)** U-ExM view of centrioles in a WT ciliated RPE1 cell stained for CEP170 (cyan), NIN (yellow) and acetylated tubulin (magenta). Bar, 200 nm. **B)** U-ExM view of centrioles in WT ciliated RPE1 cells, stained for the C-terminal domain of NIN alone, or for both the N-terminal and C-terminal domains (yellow) and for acetylated tubulin (magenta). Bar, 200 nm. **C)** NIN fluorescence intensity in the centrosome of RPE1 cells treated for 48 hours with control or NIN siRNAs then cultured for 24 hours without serum to induce ciliogenesis. Three independent experiments, N = 200 for each condition. The p value is indicated (Two-tailed T-test). **D)** Intercentriolar distance RPE1 cells treated for 48 hours with control or NIN siRNAs then cultured for 24 hours without serum to induce ciliogenesis. Three independent experiments, N > 285 for each condition. The p value is indicated (Two-tailed T-test).
